# Large scale metagenome assembly reveals novel animal-associated microbial genomes, biosynthetic gene clusters, and other genetic diversity

**DOI:** 10.1101/2020.06.05.135962

**Authors:** Nicholas D. Youngblut, Jacobo de la Cuesta-Zuluaga, Georg H. Reischer, Silke Dauser, Nathalie Schuster, Chris Walzer, Gabrielle Stalder, Andreas H. Farnleitner, Ruth E. Ley

## Abstract

Large-scale metagenome assemblies of human microbiomes have produced a vast catalogue of previously unseen microbial genomes; however, comparatively few microbial genomes derive from other vertebrates. Here, we generated 5596 metagenome-assembled genomes from the gut metagenomes of 180 predominantly wild animal species representing 5 classes, in addition to 14 existing animal gut metagenome datasets. The MAGs comprised 1522 species-level genome bins (SGBs); most of which were novel at the species, genus, or family levels, and the majority were enriched in host versus environment metagenomes. Many traits distinguished SGBs enriched in host or environmental biomes, including the number of antimicrobial resistance genes. We identified 1986 diverse biosynthetic gene clusters; only 23 clustered with any MIBiG database references. Gene-based assembly revealed tremendous gene diversity, much of it host- or environment-specific. Our MAG and gene datasets greatly expand the microbial genome repertoire and provide a broad view of microbial adaptations to the vertebrate gut.

**Importance:** Microbiome studies on a select few mammalian species (*e.g.,* humans, mice, and cattle) have revealed a great deal of novel genomic diversity in the gut microbiome. However, little is known of the microbial diversity in the gut of other vertebrates. We studied the gut microbiome of a large set of mostly wild animal species consisting of mammals, birds, reptiles, amphibians, and fish. Unfortunately, we found that existing reference databases commonly used for metagenomic analyses failed to capture the microbiome diversity among vertebrates. To increase database representation, we applied advanced metagenome assembly methods to our animal gut data and to many public gut metagenome datasets that had not been used to obtain microbial genomes. Our resulting genome and gene cluster collections comprised a great deal of novel taxonomic and genomic diversity, which we extensively characterized. Our findings substantially expand what is known of microbial genomic diversity in the vertebrate gut.

## Introduction

The vertebrate gut microbiome comprises a vast amount of genetic diversity, yet even for the most well-studied species such as humans, the number of microbial species lacking a reference genome was recently estimated to be 40-50%^1^. Uncovering this “microbial dark matter” is essential to understanding the roles of individual microbes, their intra- and inter-species diversity within and across host populations, and how each microbe interacts with each other and the host to mediate host physiology in a myriad number of ways^2^. On a more applied level, characterizing novel gut microbial diversity aids in bioprospecting of novel bioactive natural products, catalytic and carbohydrate-binding enzymes, probiotics, etc., along with aiding in the discovery and tracking of novel pathogens and antimicrobial resistance (AMR)^3^.

Recent advances in culturomic approaches have generated thousands of novel microbial genomes^4–6^, but the throughput is currently far outpaced by metagenome assembly approaches^7^. However, such large-scale metagenome assembly-based approaches have not been as extensively applied to most non-human vertebrates. The low amount of metagenome reads classified in some recent studies of the rhinoceros, chicken, cod, and cow gut/rumen microbiome suggests that databases lack much of the genomic diversity in less-studied vertebrates^8–11^. Indeed, the limited number of studies incorporating metagenome assembly hint at the extensive amounts of as-of-yet novel microbial diversity across the >66,000 vertebrate species on our planet.

Here, we developed an extensive metagenome assembly pipeline and applied it to a multi-species dataset of microbiome diversity across vertebrate species comprising 5 classes: Mammalia, Aves, Reptilia, Amphibia, and Actinopterygii, with >80% of samples obtained from wild individuals^12^ combined with data from 14 published animal gut metagenomes. Moreover, we also applied a recently developed gene-based metagenome assembly pipeline to the entire dataset in order to obtain gene-level diversity for rarer taxa that would otherwise be missed by genome-based assembly^13,14^. Our assembly approaches yielded a great deal of novel genetic diversity, which we found to be largely enriched in animals versus the environment, and to some degree, enriched in particular animal clades.

## Methods

### Sample collection

Sample collection was as described in Youngblut and colleagues^12^. Table S1A shows the dates, locations, and additional metadata of all samples collected. All fecal samples were collected in sterile sampling vials, transported to a laboratory and frozen within 8 hours. DNA extraction was performed with the PowerSoil DNA Isolation Kit (MoBio Laboratories, Carlsbad, USA).

### “multi-species” vertebrate gut metagenomes

Metagenome libraries were prepared as described by Karasov and colleagues^15^. Briefly, 1 ng of input gDNA was used for Nextera Tn5 tagmentation. A BluePippin was used to restrict fragment sizes to 400-700 bp. Barcoded samples were pooled and sequenced on an Illumina HiSeq3000 with 2×150 paired-end sequencing. Read quality control (QC) is described in the Supplemental Methods.

Post-QC reads were taxonomically profiled with Kraken2 and Bracken v.2.2^16^ against the Struo-generated GTDB-r89 Kraken2 and Bracken databases^17^. HUMAnN2 v.0.11.2^18^ was used to profile genes and pathways against the Struo-generated HUMAnN2 database created from GTDB-r89.

### Publicly available animal gut metagenomes

Published animal gut metagenome reads were downloaded from the Sequence Read Archive (SRA) between May and August of 2019. Table S1B lists all included studies. We selected studies with Illumina paired-end metagenomes from gut contents or feces. MGnify samples were downloaded from the SRA in Oct 2019 (Table S1C). Read quality control is described in the Supplemental Methods.

### Metagenome assembly of genomes pipeline

Assemblies were performed on a per-sample basis, with reads subsampled via seqtk v.1.3 to ≤20 million read pairs. The details of the assembly pipeline are described in the Supplemental Methods.

A multi-locus phylogeny of all SGB representatives was inferred with PhyloPhlAn v.0.41^19^. Secondary metabolites were identified with AntiSMASH v.5.1.1^20^ and DeepBGC v.0.1.18^21^ and then characterized with BiGSCAPE^22^. Abricate was used to identify antimicrobial resistance genes. We used Krakenuniq v.0.5.8^23^ for estimating abundance of MAGs in metagenome samples. Details can be found in the Supplemental Methods.

### Metagenome assembly of genes pipeline

Assemblies performed on a per-sample basis, with reads subsampled via seqtk v.1.3 to ≤20 million pairs. We used PLASS v.2.c7e35^14^ and Linclust (mmseqs v.10.6d92c)^13^ to assemble and cluster contigs. A full description is in the Supplemental Methods. DESeq2^24^ was used to estimate enrichment of MAGs and gene clusters in metagenomes from host and environment biomes.

### Data availability

The raw sequence data are available from the European Nucleotide Archive under the study accession number PRJEB38078. Fasta files for the 5596 non-redundant MAGs, 1522 SGBs,and gene clusters (50, 90, and 100% sequence identity clustering) can be found at http://ftp.tue.mpg.de/ebio/projects/animal_gut_metagenome_assembly/, along with genbank files for all BGCs. Code used for processing the data can be found at https://github.com/leylabmpi/animal_gut_metagenome_assembly.

## Results

### Animal gut metagenomes from a highly diverse collection of animals

We generated animal gut metagenomes from a breadth of vertebrate diversity spanning five classes: Mammalia, Aves, Reptilia, Amphibia, and Actinopterygii (the “multi-species” dataset; Figure 1). In total, 289 samples passed our read quality control, with 3.4e6 ± 5e6 s.d. paired-end reads per sample, resulting in a mean estimated coverage of 0.54 ± 0.14 s.d. (Figure S1). 180 animal species were represented, with up to 6 individuals per species (mean of 1.6). Most individuals were wild (81%).

**Figure 1.**
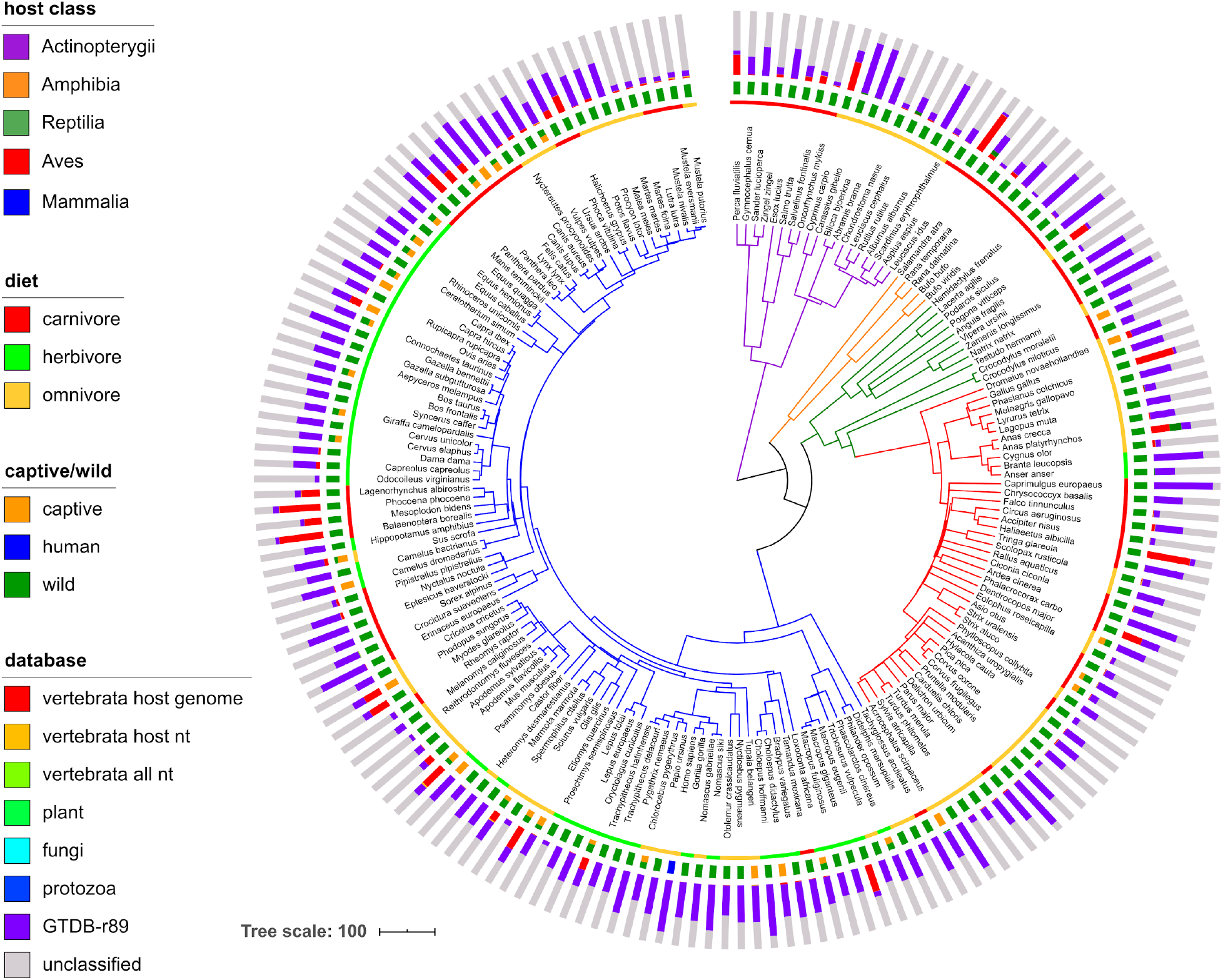
A large percentage of unmapped reads, even when using multiple comprehensive metagenome profiling databases. The dated host species phylogeny was obtained from http://timetree.org, with branches colored by host class. From inner to outer, the data mapped onto the tree is host diet, host captive/wild status, and the mean number of metagenome reads mapped to various host-specific, non-microbial, and microbial databases. Note that captive/wild status sometimes differs among individuals of the same species. The databases are i) a representative of each publicly available genome from the host species (“vertebrata host genome”), ii) all entries in the NCBI nt database with taxonomy IDs matching host species (“vertebrata host nt”), iii) as the previous, but all vertebrata sequences included, iv) the Kraken2 “plant” database, v) the Kraken2 “fungi” database, vi) the Kraken2 “protozoa” database, vii) a custom bacteria and archaea database created from the Genome Taxonomy Database, Release 89 (“GTDB-r89”). Reads were mapped iteratively to each database in the order shown in the legend (top to bottom), with only unmapped reads included in the next iteration. “unclassified” reads did not map to any database, which were used along with reads mapping to GTDB-r89 for downstream analyses (“microbial + unclassified”).

Our read-quality control pipeline included stringent filtering of host reads; some samples contained high amounts of reads mapping to vertebrate genomes (up to 74%; 6 ± 17% s.d.; Figure 1). Gut content samples contained a significantly higher amount of host reads (13.5 ± 21.6% s.d.) versus feces metagenomes (4.7 ± 12.7% s.d.; Wilcox, *P* < 1.8e-7; Table S1A). We mapped all remaining reads to a custom comprehensive Kraken2 database built from the GTDB (Release 89). Still, many samples had a low percentage of mapped reads (43 ± 22 s.d.; Figure 1), with 29% of the samples having <20% mapped reads.

### Discovery of novel diversity by large-scale metagenome assembly

Our comprehensive metagenome assembly pipeline generated 4374 non-redundant MAGs. After quality control and de-replication (see Methods), 296 MAGs remained, with a mean percent completeness and contamination of 84 ± 14 and 1.5 ± 1.2 s.d., respectively (Figure S2; Supplemental Results).

We expanded our MAG dataset by applying our assembly pipeline to 14 publically available animal gut metagenome datasets in which no MAGs have been generated by *de novo* metagenome assembly (Table S1B). Our metagenome selection included 554 samples from members of Mammalia (dogs, cats, woodrats, pigs, whales, rhinoceroses, pangolins, and non-human primates), Aves (geese, kakapos, and chickens), and Actinopterygii (cod). We applied our assembly pipeline to each individual dataset and generated a total of 5301 non-redundant MAGs (Figure S3; Supplemental Results). The substantially higher number of MAGs from these 14 datasets versus our single multi-species dataset is likely due to the larger number of samples and the high sequencing depth for many of those samples (*e.g.,* we used 2 billion paired-end reads from the dog gut microbiome dataset^25^).

We combined all MAGs and de-replicated at 99.9 and 95% average nucleotide identity (ANI) to produce 5596 non-redundant MAGs and 1522 species-level genome bins (SGBs), respectively (Tables S2A & S2B). Of the 5596 MAGs, 2773 (50%) had a completeness of ≥90%. Of the 1522 SGBs, 1184 (78%) lacked a ≥95% ANI match to the GTDB-r89, 266 (17%) lacked a genus-level match, and 6 lacked a family-level match (Figures 2 & S4). Mapping taxonomic novelty onto a multi-locus phylogeny of all 1522 SGBs revealed that novel taxa were rather dispersed across the phylogeny (Figure 2).

**Figure 2.**
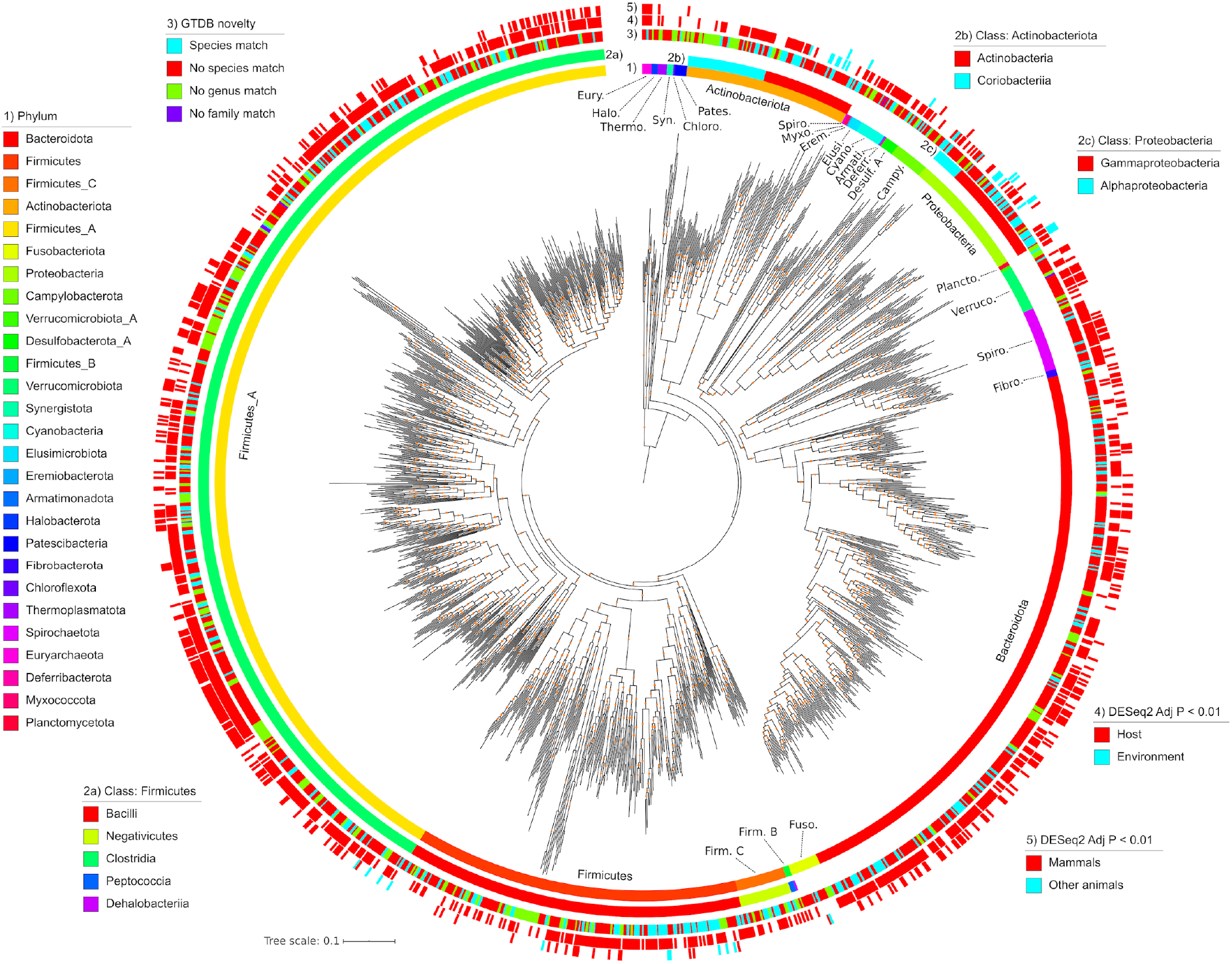
A phylogeny of all 1522 SGBs. From innermost to outermost, the data mapped onto the phylogeny is: GTDB phylum-level taxonomic classifications, class-level taxonomies for Actinobacteriota, class-level taxonomies for Firmicutes, class-level taxonomies for Proteobacteria, taxonomic novelty, significant enrichment in host gut or environmental metagenomes, and significant enrichment in Mammals or other animals in our multi-species gut metagenome dataset. The phylogeny was inferred from multiple conserved loci via PhyloPhlAn. Orange dots on the phylogeny denote bootstrap values in the range of 0.7 to 1. The phylogeny is rooted on the last common ancestor of Archaea and Bacteria.

We also assessed the novelty of our SGBs relative to UHGG, a comprehensive human gut genome database, and found that only 31% of our SGBs had ≥95% ANI to any of the 4644 UHGG representatives, and this overlap only increased to 34% at a 90% ANI cutoff.

Our SGB collection mostly consisted of MAGs assembled from a few species in the multi-study dataset, suggesting that the SGBs may not be representative of taxa found in other, more distantly related vertebrates. To assess the level of representation, we determined the prevalence of all SGBs across all multi-species metagenomes (Figure S5). The host species with the highest number of observed SGBs did tend to be those comprising the multi-study dataset (*e.g.,* pigs and primates); however, SGBs were frequently observed across the host phylogeny (41 ± 61 s.d. SGBs per host), indicating that the SGB collection was generally representative of the vertebrate gut microbiome.

Integrating the 1522 SGBs into our custom GTDB Kraken2 database significantly increased the percent reads mapped (paired t-test, *P* < 0.005; Figure S6). The percent increase varied from <1 to 62.8% (mean of 5.3 ± 6.7 s.d.) among animal species but did not appear biased to just pigs, dogs, or other vertebrate species in the multi-study datasets that we incorporated (Figure S7), which corresponds with our analysis of SGB prevalence across vertebrate hosts (Figure S5).

### Enrichment of SGBs among animal clades

While the MAGs generated here derive from animal gut metagenomes, many of these taxa might be transient in the host and actually more prevalent in the environment. We tested this by generating a “host-environment” metagenome dataset comprising 283 samples from 30 BioProjects (17 environmental and 13 host-associated; Figure 3A). We found 932 of the 1522 SGBs (61%) to be significantly enriched in the host metagenomes (DESeq2, *adj. P* < 0.01; Figure 3B). The host-enriched SGBs (host-SGBs) were taxonomically diverse, comprising 22 phyla. In contrast, only 15 SGBs (1%) were environment-enriched (env-SGBs), which all belonged to either Actinobacteriota or Proteobacteria (Figure 3B). The only SGBs that were not significantly enriched in either group belonged to Actinobacteriota or Proteobacteria, along with two SGBs from the Firmicutes A phylum. Mapping these data onto the SGB phylogeny revealed phylogenetic clustering of the environment-enriched SGBs (Figure 2).

**Figure 3.**
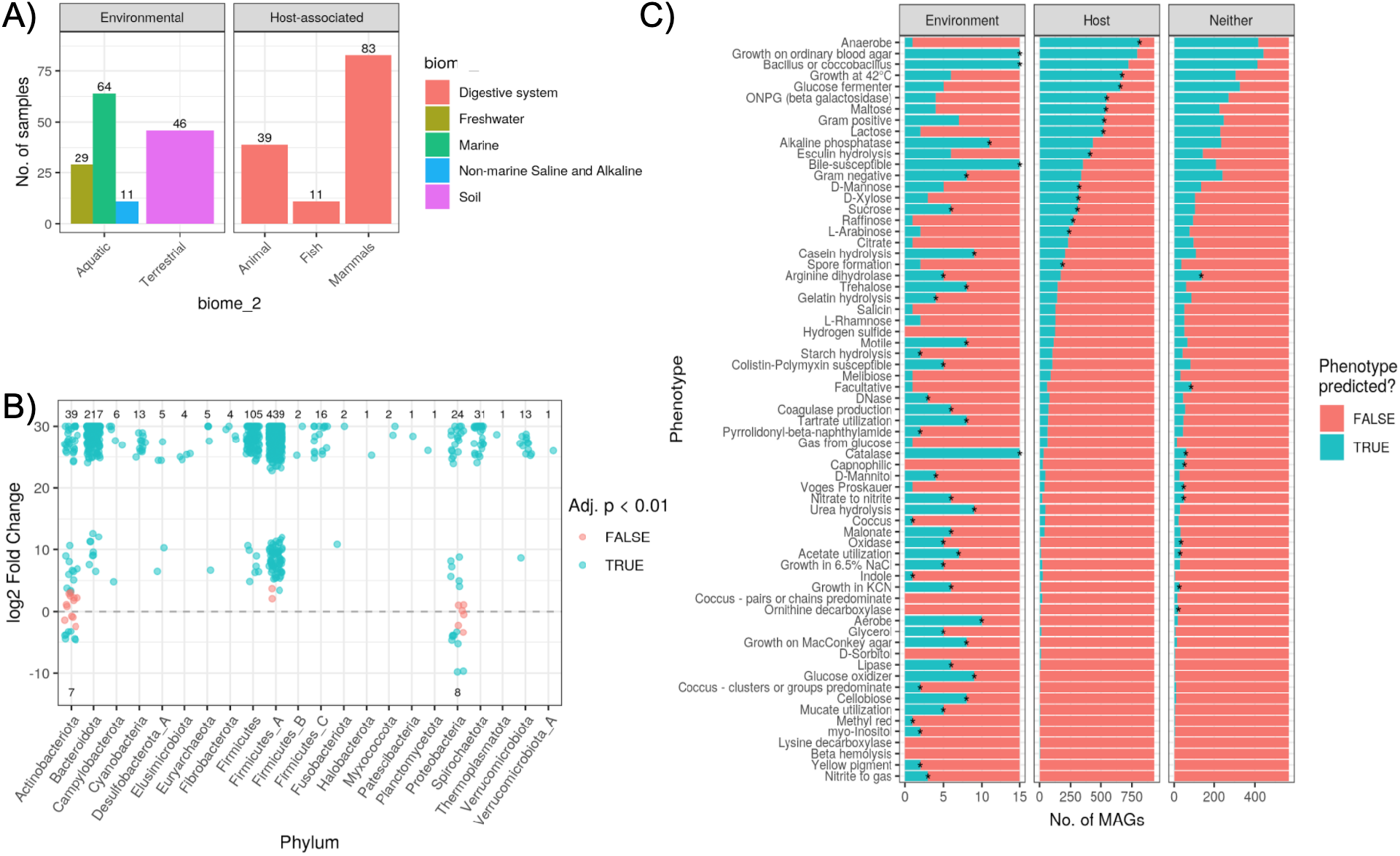
A) Summary of the number of samples per biome for our multi-environment metagenome dataset selected from the MGnify database. B) Number of SGBs found to be significantly enriched in relative abundances in host versus (positive log_2_ fold change; “l2fc”) environmental metagenomes (negative l2fc). Values shown are the number of MAGs significantly enriched (blue) in either biome or not found to be significant (red). C) Host- and environment-enriched SGBs have distinct traits. Phenotypes predicted based on MAG gene content (via Traitar ^26^) are summarized for the SGBs significantly enriched in host or environmental metagenomes (DESeq2 *Adj. P* < 0.01) or neither biome (“Neither” in the x-axis facet). Note the difference in x-axis scale. Asterisks denote phenotypes significantly more prevalent in SGBs of the particular biome versus a null model of 1000 permutations in which biome labels were shuffled among SGBs. See Table S3A for all DESeq2 results.

We investigated the traits of the host- and environment-enriched SGBs and found many predicted phenotypes to be more prevalent in one or the other group (Figure 3C; Table S2C). A total of 67 traits were predicted based on genomic content of certain pfam domains^26^. Almost all env-SGBs were aerobes (93%), which may aid in transmission between the environment and host biomes. In contrast, 87% of host-SGBs were anaerobes. Furthermore, all env-SGBs could generate catalase and were bile susceptible, while both phenotypes were sparse in host-SGBs (Figure 3C). Carbohydrate metabolism also differed, with most host-SGBs predicted to consume various tri-, di-, and mono-saccharides. In contrast, env-SGBs were enriched in phenotypes associated with motility, nitrogen metabolism, and breakdown of heterogeneous substrates (*e.g.,* cellobiose metabolism).

We also compared SGB enrichment in mammals versus non-mammals in our “multi-species” metagenome dataset and found 361 SGBs (24%) to be significantly enriched in mammals, while 22 (1%) were enriched in non-mammals (DESeq2, *adj. P* < 0.01; Figure S2C. Interestingly, 100% of SGBs in the two archaeal phyla (Halobacteria and Euryarchaeota) were enriched in mammals. Also of note, most of the Verrucomicrobiota SGBs (87%) were enriched in mammals. The only 2 phyla with >10% of SGBs enriched in non-mammals were Proteobacteria (29%) and Campylobacteria (25%).

In contrast to our assessment of phenotypes distinct to host- or env-SGBs, we did not observe such a distinction of phenotypes among SGBs enriched in Mammalia or non-mammal gut metagenomes (Figure S8). Certain phenotypes such as anaerobic growth and lactose consumption were more prevalent among mammal species, but they were not found to be significantly enriched relative to the null model.

Little is known about the distribution of antimicrobial resistance genes in the gut microbiomes of most vertebrate species^27^; therefore, we investigated the distribution of AMR genes among MAGs enriched in the environment versus host biomes. We found a mean of 35 ± 26 s.d. AMR markers per genome (Figure S9A). The high average was largely driven by Proteobacteria and Campylobacter genomes, which had a mean of 387 and 161 AMR markers per genome, respectively. The 5 most abundant markers were ruvB, galE, tupC, fabL (ygaA), and arsT (Figure S9A). The more abundant markers predominantly belonged to Firmicutes A, while Proteobacteria comprised larger fractions of the less abundant markers. Environment-enriched taxa contained substantially more AMR genes than host-enriched taxa, and the same was true for non-Mammalia versus Mammalia-enriched taxa (Figures S9B & S9C).

### MAGs reveal novel secondary metabolite diversity

We identified 1986 biosynthetic gene clusters (BGCs) among all 1522 SGBs. A total of 28 different products were predicted, with the most abundant being non-ribosomal peptide synthetases (NPRS; *n* = 473), sactipeptides (*n* = 307), and arylpolyenes (*n* = 291; Figure S10). BGCs were identified in 2 archaeal and 18 bacterial phyla. MAGs in the Firmicutes A phylum contained the most BGCs (*n* = 764; 38%), while Bacteroidota and Actinobacteriota phyla possessed 381 (19%) and 272 (14%), respectively (Figure S10). Still, Actinobacteriota SGBs did possess the highest average number of BGCs per genome (16.3), followed by Eremiobacterota (9), Proteobacteria (7.7), and Halobacterota (5.1).

Clustering all 1986 BGCs by BiGSCAPE generated 1764 families and 1305 clans, with clans being a second, coarser level of clustering^22^. Only 8 clans (comprising 23 BGCs) included any MIBiG database reference, suggesting a high degree of novelty (Figure S11). Mapping the BGCs on a genome phylogeny of all species containing ≥3 BGCs (233 SGBs) revealed that the number of BGCs per genome was somewhat phylogenetically clustered: the five genomes with the most BGCs belonged either to the Actinobacteria or Gammaproteobacteria (Figure 4). Notably, these clades contained a high number of host-SGBs. Of these 233 SGBs, the majority were taxonomically novel, with 62% lacking a species-level match to GTDB-r89, and 18% lacking a genus-level match (Figure 4). To determine which of the BGCs are most prevalent across animal hosts, we quantified the prevalence of each BGC family across our multi-species metagenome dataset and mapped it to the genome phylogeny (Figures 4 & S12). Of the 1543 BGC families found in the 233 SGBs, 83 were present in ≥25% of the animal metagenomes, with ribosomally synthesized and post-translationally modified peptides (RIPPs) being by far the most prevalent (up to 98% prevalence of individual BGC families) and also found in species from a number of phyla.

**Figure 4.**
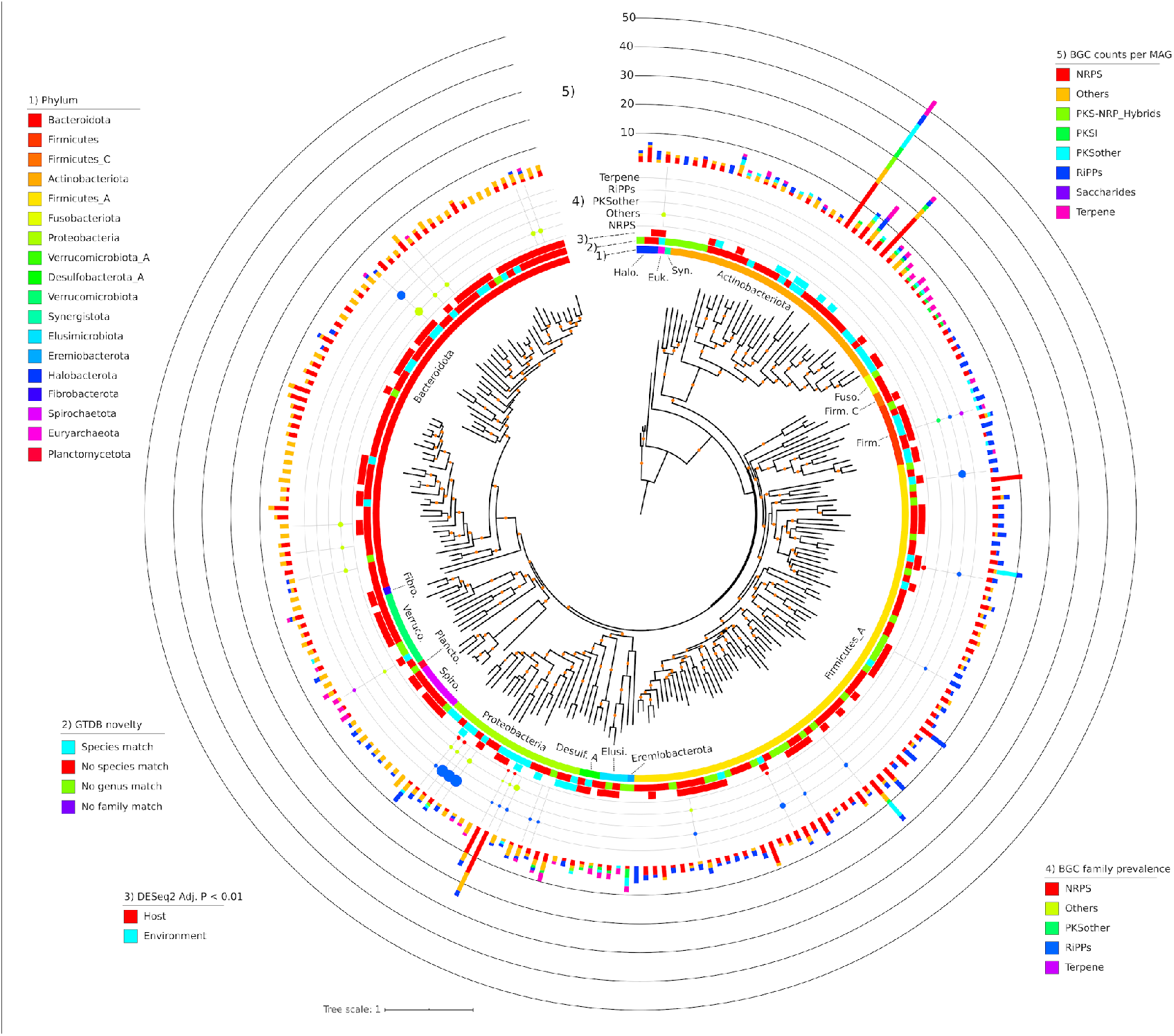
Phylogeny of all SGBs (*n* = 233) with ≥3 BGCs identified by AntiSMASH. From innermost to outermost, the data mapped onto the phylogeny is: 1) GTDB phylum-level taxonomic classifications, 2) taxonomic novelty, 3) significant enrichment in host or environmental metagenomes, 4) the prevalence of BGC families across the multi-species metagenome dataset, and 5) the number of BGCs identified in the MAG. Prevalence is the maximum of any BGC family for that BGC type, and only BGC families with a prevalence of ≥25% are shown. The phylogeny is a pruned version of that shown in Figure 2. Orange dots on the phylogeny denote bootstrap values in the range of 0.7 to 1.

### Large-scale gene-based metagenome assembly reveals novel diversity

We applied gene-based assembly methods to our combined metagenome dataset^14^, which generated a total of 150,718,125 non-redundant coding sequences (average length of 179 amino acids). Clustering at 90 and 50% sequence identity resulted in 140,225,322 and 6,391,861 clusters, respectively. Only 16.9 and 11.3% of each respective cluster set mapped to the UniRef50 database, indicating that most coding sequences were novel. The clusters comprised 88 bacterial and 11 archaeal phyla; 80 of which were represented by <100 clusters, and 60 lacking a cultured representative. Proteobacteria (mostly Gammaproteobacteria), Firmicutes, and Bacteroidetes made up 92.2% of all clusters (Figure 5A). The proportion of clusters belonging to each COG functional category was largely the same for the more abundant bacterial phyla (Figure 5B), while more variation was seen among Euryarchaeota (Figure 5C). The dominant 7 phyla showed substantial variation in the number of clusters associated with various KEGG pathway categories (Figure S14). For instance, a high proportion of Fusobacteria and Tenericutes clusters were associated with the “nucleotide metabolism”, “replication and repair”, and “translation” categories. A total of 87,573 clusters were annotated as CAZy families, with GT51, GH13, GH18, GT02, and GT04 representing 48% of all CAZy-annotated clusters (Figure 5E). Of the 12 phyla with the most CAZy family clusters, there were substantial differences in proportions of clusters falling into each family (Figure 5F).

**Figure 5.**
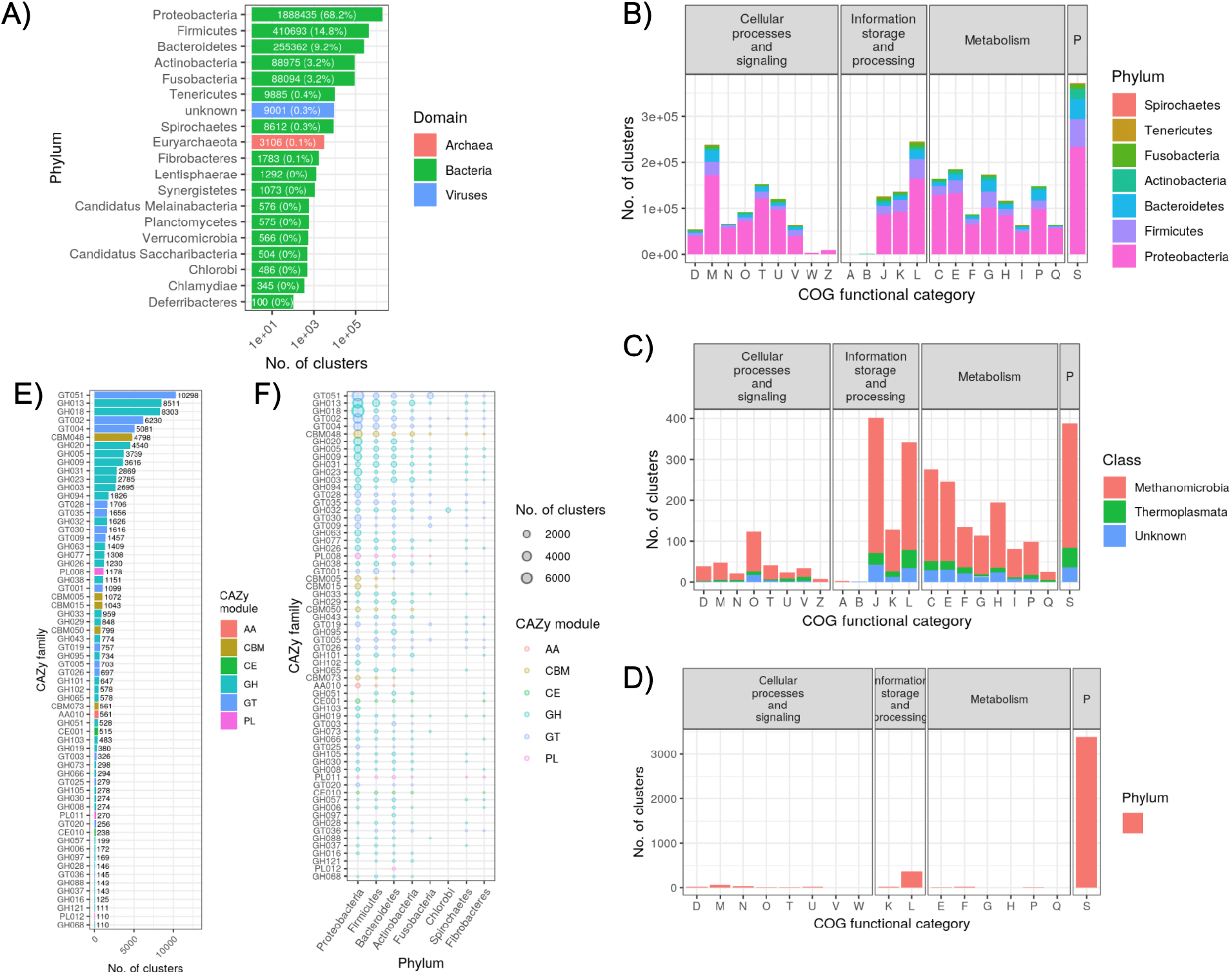
A summary of the 50% sequence identity clusters generated from the gene-based metagenome assembly of the combined dataset. A) The total number of gene clusters per phylum. For clarity, only phyla with ≥100 clusters are shown. Labels on each bar list the number of clusters (and percent of the total). B) The number of bacterial gene clusters per phylum and COG category. The “P” facet label refers to “poorly characterized”. C) The number of archaeal gene clusters per class (all belonging to Euryarchaeota) and COG category. D) The number of viral gene clusters per COG category. E) The number clusters annotated as each CAZy family. For clarity, only phyla with ≥100 clusters are shown. Labels next to each bar denote the number of clusters. F) The number of clusters per CAZy family, broken down by phylum. CAZy families and phyla are ordered by most to least number of clusters. For clarity, only CAZy families and phyla with ≥100 total clusters are shown.

### Biome enrichment of gene clusters from specific phyla

We mapped reads from our host-environment metagenome dataset to each cluster and used DESeq2 to identify those significantly enriched (*adj. P* < 1e-5) in each biome. Most strikingly, the same functional groups were enriched in both biomes, regardless of the grouping (*i.e.,* COG functional category, KEGG pathway, or CAZy family); however, the gene clusters belonged to different microbial phyla (Figure 6; Supplemental Results). For instance, nearly all COG categories for gene clusters belonging to Proteobacteria were environment-enriched, while the same COG categories for clusters belonging to Firmicutes and Bacteroidetes were host-enriched. In contrast, functional groups of certain phyla were enriched in one biome, while different groups were enriched in the other, indicating within-phylum differences in functional content and habitat distributions. For instance, Fusobacteria KEGG pathways were predominantly host-enriched, but protein export, bacteria secretion system, and aminoacyl-tRNA biosynthesis were environment-enriched, indicating that these 3 pathways were more predominant in environment-enriched members of Fusobacteria (Figure 6B). Overall, these results suggest that both biomes select for these same microbial functions, but the microbes involved often differ at coarse taxonomic scales.

**Figure 6.**
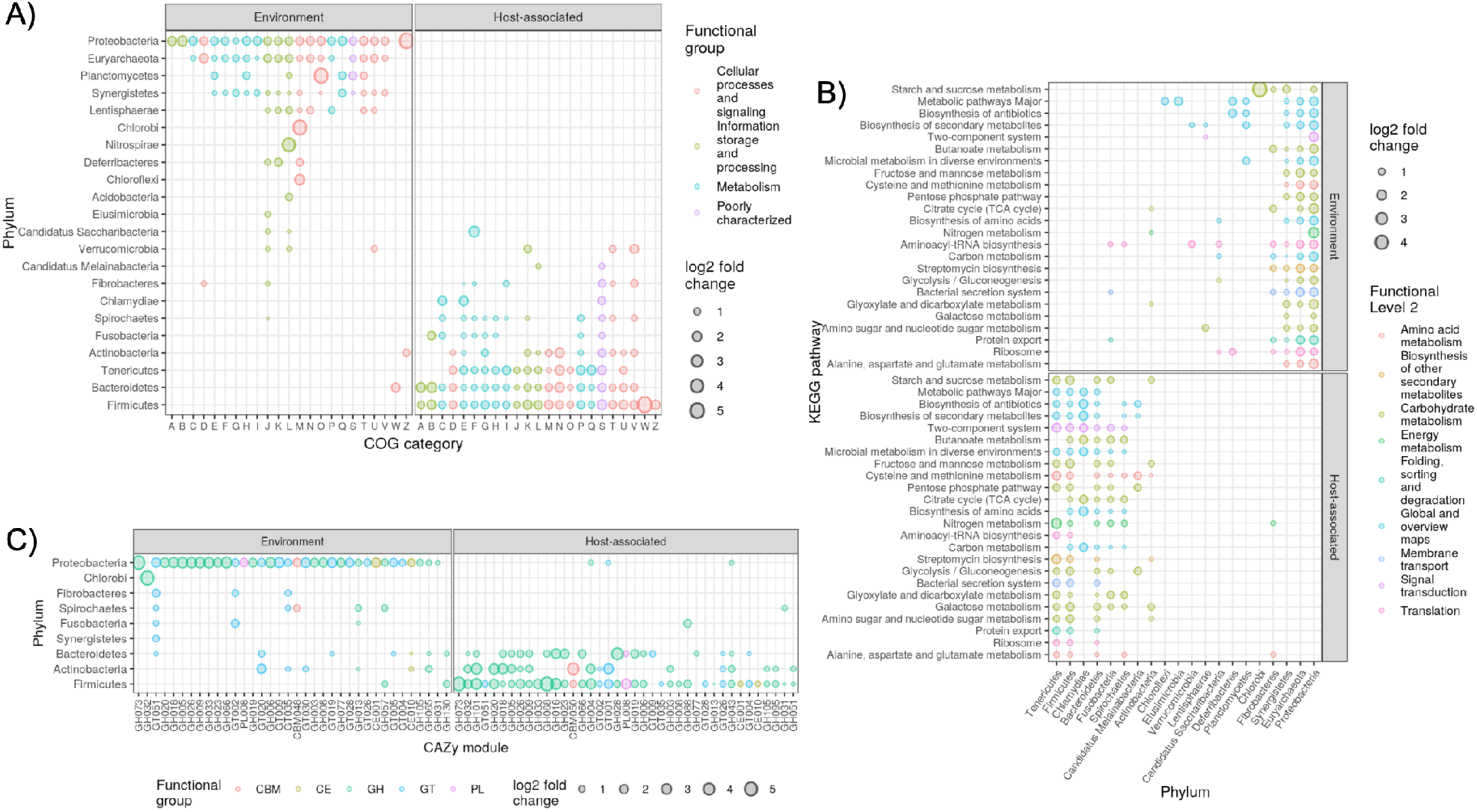
Enrichment of gene clusters grouped by phylum and A) COG category B) KEGG pathway or C) CAZy family. Only groupings significantly enriched in abundance (DESeq2, *adj. P* < 1e-5) in either biome are shown. Only gene clusters observed in at least 25% of the metagenomes were included. For clarity, only KEGG pathways enriched in >7 phyla are shown, and only CAZy families enriched in >1 phylum are shown. Note that the axes are flipped in B) relative to A) and C). See Tables S5A, S5B, and S5C for all DESeq2 results.

We also assessed gene cluster enrichment in Mammalia versus non-Mammalia and found fewer significantly enriched features, which may be due to the smaller metagenome sample size or less pronounced partitioning of functional groups among biomes (Figure S15; Supplemental Results). Still, we again observed that both biomes enriched for the same microbial functions, but these belonged to different coarse taxonomic groups. To assess whether abundance estimations were substantially erroneous due to mis-mapping of metagenome reads to the gene clusters, we reran the analysis with stricter DIAMOND mapping parameters but observed similar findings, even though 48% fewer gene clusters were detected in any metagenome (Figure S16).

### Functional metagenome profiling benefits from our gene catalogue

Lastly, we integrated our gene catalogue into a custom HUMAnN2 database built from the GTDB-r89 and found that this combined database substantially increased mappability of reads from our multi-species metagenome dataset (Figure S17; Supplemental Results).

## Discussion

Our MAG and gene cluster datasets, derived from 289 newly generated metagenomes from 180 vertebrate species, along with 544 metagenomes from 14 publicly available animal gut metagenome datasets, substantially helps to expand the breadth of cross-species gut metagenome comparisons (Figures 1 & 5). While metagenomics is rapidly expanding in popularity^7^, most analyses of metagenomic data suffer from a reliance on incomplete reference databases^17^, which we show to be acutely problematic for the gut microbiomes of most vertebrates in our dataset (Figure 1). Grossly incomplete surveys of microbial diversity can lead to incorrect findings on community assembly in the vertebrate gut^28^. Although our dataset has only partially revealed this unknown diversity, it does substantially improve reference database coverage at both the genome and gene levels and also provides an estimate of the incompleteness of existing reference databases.

A major contribution of this study is the extensive MAG collection that we generated by assembling the metagenomes of our multi-species dataset together with 14 other animal gut metagenome datasets from understudied host species. This collection includes 1184, 266, and 6 genomes from novel species, genera, and families, respectively (Figures 2 & S4). Moreover, we found little overlap (31%) between our MAG collection and the extensive human microbiome genome catalogue comprising the UHGG, which underscores its taxonomic novelty. We also showed substantial SGB prevalence across all 5 vertebrate taxonomic classes (Figure S5), indicating that our MAG collection is representative microbes found across the vertebrate taxonomy. Our MAG collection, once combined with the GTDB^29^, improved our ability to classify reads in our multi-species metagenome dataset (Figure S6), which is critical for accurately assessing gut microbiome diversity across vertebrates. Although MAGs have been criticized for their incompleteness and potentially high prevalence of misassemblies^30^, we note that i) the overall completeness of our MAGs was rather high (90% median completeness), ii) complete genomes are not required for accurate taxonomic profiling^31^, iii) the prevalence of misassemblies among MAGs is likely quite low when using state-of-the-art assembly and binning approaches^32^. Still, researchers who may utilize this set of MAGs should use caution when analyzing individual SNPs, plasmids, genomic islands, or other potentially missing or misassembled genomic features^33^.

We investigated the distribution of our MAGs across environment and host biomes to elucidate the diversity of host-microbe symbiosis in the vertebrate gut. Microbe-host symbiosis spans the continuum from free-living microbes that can simply survive passage through the host gut, to obligate symbioses^34^. Therefore, MAGs enriched in the environment versus the host would indicate a weak association, while the opposite enrichment would suggest a more obligate symbiosis. We provide evidence of host specificity for the majority of SGBs, while a few Proteobacteria and Actinobacteria SGBs were environment-enriched. When just considering host-associated metagenomes, these env-SGBs were generally enriched in non-mammals (Figures 2, 3, & S8). This is consistent with the hypothesis that mixed-mode transmission, especially between environmental sources and hosts, is more commonplace in non-mammalian gut microbiome community assembly versus in mammals^35^.

Our trait-based analysis of SGBs supports the notion that host-enriched taxa are adapted for a symbiotic lifestyle, while environment-enriched taxa are adapted for a free-living or facultative symbiosis lifestyle (Figure 3). For instance, anaerobes comprised almost all host-enriched SGBs, while environment-enriched SGBs were aerobes or facultative anaerobes and generally motile, which could be highly beneficial for transmission between the environment and gut biomes. Indeed, a recent directed evolution experiment showed that selecting for inter-host migration can generate bacterial strains with increased motility^36^, and a trait-based study of the human infant gut microbiome showed that later stages of succession are dominated by taxa adapted to the anoxic gut^37^.

By assessing SGB enrichment in Mammalia versus non-Mammlia metagenomes, we elucidated the specificity of host-microbe symbioses in the gut across large evolutionary distances. More SGBs were enriched in mammals versus non-mammals (Figures 2 & S8), as we observed in our previous 16S rRNA assessment of these vertebrate clades^12^. Few traits differed among SGBs enriched in either biome (Figure S8), which may indicate that the traits assessed are similarly required for adaptation to each host clade, even at this coarse evolutionary scale.

Vertebrates both play a critical role in the spread of antimicrobial resistance and also have been sources of novel antibiotics and other natural products^27,38^. We investigated BGC and AMR diversity in our MAG collection and observed a high diversity of BGC products, but very few of the BGCs clustered into families with experimentally characterized BGCs from the MIBiG database (Figures S9 & S10). This contrasts with findings that only ~10% of BGCs in the human microbiome are uncharacterized^39^, which is likely due to the limited study of natural products in the gut microbiome of non-human vertebrates^40,41^. We found NRPS-producing BGCs to be prevalent among the Firmicutes SGBs, which is similar to a recent assessment of 501 genomes from rumen isolates in which thousands of BGCs were identified^42^. Still, RiPPs were most prevalent across all vertebrate clades, which expands upon observations of high prevalence of this BGC class in the gut microbiome of humans^39^ (Figures 2 & 4).

By combining our AMR marker screen with our SGB biome enrichment analysis, we were able to characterize how AMR is associated with varying degrees of symbiosis (Figure S9), which is important for understanding AMR reservoirs^27,43^. Our findings indicate that the AMR reservoir may be greater for free-living and facultatively symbiotic taxa relative to microbes with stronger host associations (Figure S9). Indeed, some of the most abundant AMR markers were associated with metal resistance (*e.g.,* ruvB, tupC, and arsT), which may reflect a lifestyle in which the microbe is exposed to environmental sources of metals^44,45^.

While MAGs provide a powerful means of investigating species and strain-level diversity within the vertebrate gut microbiome, the approach is limited to only relatively abundant taxa with enough coverage to reach adequate assembly contiguity^46^. Our gene-based assembly approach allowed us to greatly expand the known gene catalogue of the vertebrate gut microbiome beyond just the abundant taxa, with a total of >150 million non-redundant coding sequences generated, comprising 88 bacterial and 11 archaeal phyla (Figure 5). In comparison, recent large-scale metagenome assemblies of the gut microbiome from chickens, pigs, rats, and dogs have generated 7.7, 9.04, 7.7, 5.1, and 1.25 million non-redundant coding sequences, respectively^8,25,47,48^. It is also illustrative to consider that a recent large-scale metagenome assembly of cattle rumen metagenomes generated 69,678 non-redundant genes involved in carbohydrate metabolism^9^, while our gene collection comprised substantially more CAZy-annotated gene clusters (*n* = 87,573), even after collapsing at 50% sequence identity. The increased mappability that we achieved across all 5 vertebrate clades when incorporating our gene catalogue in our functional metagenome profiling pipeline demonstrates how our gene collection will likely aid future vertebrate gut metagenome studies (Figure S17).

Our assessment of gene cluster abundances in metagenomes from environment and host-associated biomes illuminates how microbiome functioning and taxonomy is distributed across the free-living to obligate symbiont spectrum. Most notably, nearly all prominent functional groups were enriched in both the environment and host-associated biomes, but the specific gene clusters belonged to different taxonomic groups in each biome (Figure 6). For instance, almost all abundant CAZy families were enriched in both the environment and host biomes, but the environment was dominated by Proteobacteria, while Firmicutes, Bacteroidetes, and Actinobacteria gene clusters comprised most host-enriched CAZy families. This suggests the same coarse-level functional groups are present across the free-living to obligate microbe-vertebrate symbiosis lifestyles, but coarse-level taxonomy strongly differs across this spectrum. This pattern largely remained true when we compared enrichment between the Mammalia and non-mammals, suggesting that taxonomic differences prevail over functional differences in regards to host specificity, at least over broad-scale vertebrate evolutionary distances. While comparing function to taxonomy is challenging due to differing levels of resolution, we do not believe that our findings are simply due to using functional groupings that are coarser than taxonomy, given that i) we assessed multiple functional grouping (COG, KEGG, and CAZy), which all showed similar patterns, even though they differ in functional resolution, and ii) we assessed taxonomy at the very coarse phylum level but still found stark taxonomic differences across biomes.

In conclusion, our large-scale metagenome assembly of both MAGs and coding sequences from a diverse collection of vertebrates substantially expands the known taxonomic and functional diversity of the vertebrate gut microbiome. We have demonstrated that both taxonomic and functional metagenome profiling of the vertebrate gut is improved by our MAG and gene catalogues, which will aid future investigations of the vertebrate gut microbiome. Moreover, our collection can help guide natural product discovery and bioprospecting of novel carbohydrate-active enzymes, along with modeling AMR transmission among reservoirs. By characterizing the distribution of MAGs and microbial genes across environment and host biomes, we gained insight into how taxonomy and function differ along the free-living to obligate symbiosis lifestyle spectrum. We must note that our metagenome assembly dataset is biased toward certain animal clades, which likely impacts these findings. As metagenome assembly becomes more commonplace for studying the vertebrate gut microbiome, bias toward certain vertebrates (*e.g.,* humans) will decrease, and thus allow for a more comprehensive reassessment of our findings.

## Acknowledgements

We thank Nadine Ziemert for helpful discussions in regards to bioinformatic approaches for secondary metabolite detection and analysis. This work was supported by the Department of Microbiome Science at the Max Planck Institute for Developmental Biology. This study was supported by the Austrian Science Fund (FWF) research projects P23900 granted to Andreas H. Farnleitner and P22032 granted to Georg H. Reischer. Further support came from the Science Call 2015 “Resource und Lebensgrundlage Wasser” Project SC15-016 funded by the Niederösterreichische Forschungs- und Bildungsgesellschaft (NFB).

We would like to thank the following collaborators for their huge efforts in sample and data collection: Mario Baldi, School of Veterinary Medicine, Universidad Nacional de Costa Rica; Wolfgang Vogl and Frank Radon, Konrad Lorenz Institute of Ethology and Biological Station Illmitz; Endre Sós and Viktor Molnár, Budapest Zoo; Ulrike Streicher, Conservation and Wildlife Management Consultant, Vietnam; Katharina Mahr, Konrad Lorenz Institute of Ethology, University of Veterinary Medicine Vienna and Flinders University Adelaide, South Australia; Peggy Rismiller, Pelican Lagoon Research Centre, Australia; Rob Deaville, Institute of Zoology, Zoological Society of London; Alex Lécu, Muséum National d’Histoire Naturelle and Paris Zoo; Danny Govender and Emily Lane, South African National Parks, Sanparks; Fritz Reimoser, Research Institute of Wildlife Ecology, University of Veterinary Medicine Vienna; Anna Kübber-Heiss and Team, Pathology, Research Institute of Wildlife Ecology, University of Veterinary Medicine Vienna; Nikolaus Eisank, Nationalpark Hohe Tauern, Kärnten; Attila Hettyey and Yoshan Moodley, Konrad Lorenz Institute of Ethology, University of Veterinary Medicine Vienna; Mansour El-Matbouli and Oskar Schachner, Clinical Unit of Fish Medicine, University of Veterinary Medicine; Barbara Richter, Institute of Pathology and Forensic Veterinary Medicine, University of Veterinary Medicine Vienna; Hanna Vielgrader and Zoovet Team, Schönbrunn Zoo; Reinhard Pichler, Herberstein Zoo. We explicitly thank the Freek Venter of South African National Parks and the National Zoological Gardens of South Africa for granting access to their Parks for sample collection.

## Author Contributions

G.H.R., R.E.L., and A.H.F. created the study concept. G.H.R., N.S., C.W., and G.S. performed the sample collection and metadata compilation. G.H.R., N.S., and S.D. performed the laboratory work. N.D.Y. and J.C. performed the data analysis. N.D.Y., J.C., and R.E.L. wrote the manuscript.

