## Supplemental Materials for "Large scale metagenome assembly reveals novel animal-associated microbial genomes, biosynthetic gene clusters, and other genetic diversity"

### Supplemental Methods

*"multi-species" metagenomes*

Metagenome libraries were prepared as described by Karasov and colleagues (Karasov et al. 2018). Briefly, 1 ng of input gDNA was used for Nextera Tn5 tagmentation. A BluePippin was used to restrict fragment sizes to 400-700 bp. Barcoded samples were pooled and sequenced on an Illumina HiSeq3000 with 2x150 paired-end sequencing.

Raw reads were validated with fqtools v.2.0 (Droop 2016) and de-duplicated with the bbtools v.37.78 "clumpify" command (https://jgi.doe.gov/data-and-tools/bbtools/). Skewer v.0.2.2 (Jiang et al. 2014) and bbtools "bbduk" command were used for adapter trimming and sequence quality control filtering. We used the bbtools "bbmap" command to filter reads mapping to the human genome hg19 assembly. Read quality reports for each step were generated with fastqc v.0.11.7 (https://github.com/s-andrews/FastQC) and multiQC v.1.5a (Ewels et al. 2016).

We filtered out host reads from the metagenomes through iterative filtering across multiple databases with Kraken2 v.2.0.8 (Parks et al. 2018). The databases and order of filtering are listed in Figure 1. Host genomes were downloaded from NCBI in June 2019 (see Table S1D for a list of genome assembly accessions). blastcmd was used to obtain entries in the NCBI nt database with taxonomy IDs matching host species. Low complexity sequences were removed with bbduk.sh (-Xmx100g threads=24 entropy=0.1 entropywindow=50 entropyk=5). Struo v.0.1.6 (de la Cuesta-Zuluaga, Ley, and Youngblut 2019) was used to build a custom Kraken2 Bacteria and Archaea database created from the Genome Taxonomy Database (GTDB), Release 89 (Parks et al. 2018).

After read filtering, coverage of community diversity was estimated with nonpareil v.3.3.3 (Rodriguez-R et al. 2018). Estimated coverage of taxonomic diversity did not correlate with number of reads (Figure S1), suggesting that the microbiome diversity is highly variable among host species, which corresponds with our previous work (Youngblut et al. 2019). We therefore filtered metagenome samples with <30% estimated coverage and not based on number of post-QC reads.

Filtered reads were taxonomically profiled with Kraken2 and Bracken v.2.2 (Lu et al. 2017) against the Struo-generated GTDB-r89 Kraken2 and Bracken databases (de la Cuesta-Zuluaga, Ley, and Youngblut 2019). Humann2 v.0.11.2 (Franzosa et al. 2018) was used to profile genes and pathways against the Struo-generated HUMAnN2 database created from GTDB-r89.

*Publicly available animal gut metagenomes*

Published animal gut metagenome reads were downloaded from the Sequence Read Archive (SRA) between May and August of 2019. Table S1B lists all included studies. We selected studies with Illumina paired-end metagenomes from gut contents or feces. MGnify (Mitchell et al. 2020) samples were downloaded from the SRA in Oct 2019 (Table S1C). Read quality control for published metagenomes was the same as for the metagenomes generated in this work, except "clumpify" from bbtools was not used due a lack of flow cell location information in the sequence headers.

*Metagenome assembly of genomes pipeline*

Assemblies were performed on a per-sample basis, with reads subsampled via seqtk v.1.3 to at most 20 million read pairs. In order to reduce the computational load of *de novo* assembly, we first performed reference-based metagenome assembly on each sample with metacompass v.1.2 (Cepeda et al. 2017), which itself used Pilon v.1.22 (Walker et al. 2014) to create reference-based contigs. To select reference genomes, we profiled all metagenomes via Kraken2 and Bracken against our custom GTDB-r89 database, and the genomes of all species with a relative abundance of ≥1% were downloaded via ncbi-genome-download v.0.2.1 based on species' NCBI taxIDs (RefSeq). dRep v.2.4 was then used to de-replicate the reference genomes to 99% ANI representatives. All reads not mapping to references and thus used for the reference-based assemblies were used for per-sample *de novo* assemblies via metaSPAdes v.3.12.0 (Nurk et al. 2017) , which has been shown to generally be more accurate than Megahit (Vollmers, Wiegand, and Kaster 2017) but requires more computational resources. Contigs of ≥2000 bp from the reference-based and *de novo* assemblies were combined and de-replicated with the bbtools "dedup.sh" script (minidentity=100 minscaf=500 minoverlappercent=95).

Contig binning was also per-sample, but reads from all metagenomes were used for calculating differential coverage in each binning approach. Two binners were used: MaxBin2 v.2.2.4 (Wu, Simmons, & Singer, 2016) and MetaBAT2 v.2.12.1 (Kang et al., 2019), with 2 parameter settings each for a total of 4 bin collections per sample. For MaxBin2, the parameter settings were "-min_contig_length 2000 -markerset 40 -prob_threshold 0.6" and "-min_contig_length 2000 -markerset 40 -prob_threshold 0.8". For MetaBAT2, the parameter settings were "–minContig 2000 –minCV 0.5 –minCVSum 0.5 –maxP 92 –maxEdges 150" and "–minContig 2000 –minCV 0.5 –minCVSum 0.5 –maxP 97 –maxEdges 500". All binning methods incorporated differential coverage (coverage of each contig in each metagenome), which has not been widely employed for large scale metagenome assemblies due to its high computational burden but can generate higher quality assemblies (Meyer et al. 2018). Coverage was determined by mapping reads from all metagenomes within each study to each per-sample assembly contigs via Bowtie2 v.2.3.5 (Langmead and Salzberg

2012). DAS-Tool v.1.1.1 (Sieber et al. 2018) was used to select the highest quality non-redundant set of bins (MAGs) from all binning runs, with quality based on CheckM-estimated contamination and completeness (Parks et al. 2015). We found that DAS-Tool selected some MAGs from each binning approach.

MAGs selected by DAS-Tool from each per-sample metagenome were combined across all samples. MAG contamination and completeness were assessed via CheckM v.1.0.13, and MAG taxonomy was assessed via CheckM, Sourmash v.2.3.0 (Pierce et al. 2019) and GTDB-Tk v.0.3.3 (Chaumeil et al. 2019). MAG pairwise ANI was calculated with fastANI v.1.2 (Jain et al. 2018). All MAGs with a CheckM-estimated completeness of <50% or contamination of ≥5% were removed from downstream analyses. MAGs were de-replicated at 99.9% ANI with dRep to collapse clonal genomes while accounting ANI differences due to misassemblies. MAGs were de-replicated with dRep to 95% ANI to create species-level genome bins (SGBs) (Olm et al. 2020).

A multi-locus phylogeny of all SGB representatives was inferred with PhyloPhlAn v.0.41 (Segata et al. 2013). Secondary metabolites were identified with AntiSMASH v.5.1.1 (Blin et al. 2019) and DeepBGC v.0.1.18 (Hannigan et al. 2019). The number of BGCs identified by DeepBGC differed greatly depending on the confidence cutoff but generally matched AntiSMASH in the cutoff range of 0.7-0.8. We used BiGSCAPE (Navarro-Muñoz et al. 2020) for clustering BGCs and comparing them to known BGCs in the MIBiG database (Kautsar et al. 2020). Abricate was used to identify antimicrobial resistance genes (parameters: “--minid 75 --mincov 80”), with the BLAST hits to the following databases aggregated: NCBI AMRFinderPlus, CARD, Resfinder, ARG-ANNOT, BacMet2, VFDB, MEGARES2, PlasmidFinder. We used Krakenuniq v.0.5.8 (Breitwieser, Baker, and Salzberg 2018) for estimating abundance of MAGs in metagenome samples, with a unique kmer cutoff of >1000.

*Metagenome assembly of genes pipeline*

Assemblies performed on a per-sample basis, with reads subsampled via seqtk v.1.3 to at most 20 million pairs. PLASS v.2.c7e35 (Steinegger, Mirdita, and Söding 2019) was used for gene-based assembly at a minimum sequence identity of 0.9. We used Linclust (mmseqs v.10.6d92c) (Steinegger and Söding 2018) to cluster all genes at three minimum sequence identity cutoffs: 100, 90, and 50%. Genes were annotated with eggnog-mapper v.2.0.1 (Huerta-Cepas et al. 2017) against the eggNOG 5 database (Huerta-Cepas et al. 2019). We used two methods taxonomically classify genes: i) the mmseqs “taxonomy” subcommand with the UniClust50 2018-08 database, and ii) DIAMOND v.0.9.28 (“blastp --evalue 1e-5 --top 10 --sensitive”) against the NCBI nr database. We found DIAMOND to be more accurate and sensitive than the mmseqs2 taxonomic classification workflow, although more computationally intensive (Figure S13). Specifically, DIAMOND classified more gene clusters, and the taxonomic classifications were more reasonable, given that only mmseq2 generated a long “tail” of candidatus phyla with few clusters per phylum. DIAMOND (“blastx --evalue 0.001 --sensitive --max-target-seqs 10”) was used to estimate gene cluster abundances in metagenomes. We assessed mapping sensitivity by performing the same analysis with stricter mapping parameters (“blastx --id 50 --subject-cover 0.8 --evalue 0.001 --sensitive --max-target-seqs 10”; Figure S16). Any clusters with <80% coverage across the sequence were set to zero abundances. DESeq2 (Love, Huber, and Anders 2014) was used to estimate enrichment of relative abundances in metagenomes from host and environment biomes.

*General data analysis*

Read quality control, metagenome profiling, metagenome assembly of genomes, and metagenome assembly of genes were all run as pipelines on a computer cluster with Snakemake v.5.4.5 (Köster and Rahmann 2012). All general data processing was performed in R (R Core Team 2020). The dplyr (Wickham and Francois 2016), tidyr (Wickham 2016), ggplot2 (Wickham 2009), readxl (Wickham and Bryan 2019), data.table (Dowle et al. 2014), tidytable (Fairbanks 2020), future (Bengtsson 2019b), and future.batchtools (Bengtsson 2019a) R packages were used for general data processing and figure generation. The non-default software parameters used for the analyses in this study are listed in Table S1E.

### Supplemental Results

*Multi-species dataset: MAG taxonomy*

The 296 MAGs generated from the multi-species metagenome dataset consisted of 11 bacterial and 1 archaeal phylum, as determined via GTDB-Tk (Chaumeil et al. 2019). The majority of MAGs belonged to the classes Clostridia ($n$ = 95; Firmicutes A phylum) and Bacteroidia ($n$ = 74; Bacteroidota phylum; Figure S2). De-replicating MAGs at 95% ANI produced 248 species-level genome bins (SGBs). Of the SGBs, 196 (79%) had <95% ANI to every genome in the GTDB-r89 database, and 51 (21%) lacked a genus-level match. These findings indicated that the MAG dataset contained a substantial amount of novel diversity.

*MAGs generated from published animal gut metagenomes*

As with the multi-species metagenome assemblies, MAG quality was high, with a mean completeness and contamination of 85 ± 13 and 1.1 ± 1.1 s.d., respectively. The taxonomic diversity was also quite high, with 2 archaeal and 25 bacterial phyla represented (Figure S3). De-replicating MAGs at 95% ANI produced 1308 SGBs. Of these, 1001 lacked a ≥95% ANI match to the GTDB-r89, 216 lacked a genus-level match, and 6 lacked even a family-level match

*Environment versus host enrichment of gene clusters*

For COG categories, Proteobacteria, Euryarchaeota, Planctomycetes, Synergistetes, and Lentisphaerae were strongly environment-enriched, while Firmicutes, Bacteroidetes, Tenericutes, Actinobacteria, Fusobacteria, and Spirochaetes were strongly host-enriched (Figure 6A). In contrast, some phyla comprised a mixture of environment- and host-enriched categories. For instance, Verrucomicrobia-classified clusters in the J (Translation, ribosomal structure and biogenesis), L, and U (Intracellular trafficking, secretion, and vesicular transport) categories were enrichment-enriched, while the K (Transcription), T (Signal transduction mechanisms), and V (Defense mechanisms) categories were host-enriched, which likely reflects broad-scale intra-phylum adaptive differentiation to each environment (Figure 6A). Interestingly, the W (Extracellular structures) and Z (Cytoskeleton) categories were environment-enriched for Bacteroidetes and Actinobacteria, respectively, while most other categories for these phyla were host-enriched, suggesting that these specific functional groups are of increased benefit for environment-predominant members of the clades.

A similar pattern was observed when assessing the distribution of KEGG pathways. Nearly all dominant pathways for Proteobacteria, Euryarchaeota, and Synergistetes environment-enriched, while nearly all pathways for Firmicutes and Tenericutes were host-enriched (Figure 6B). In contrast, some pathways of certain phyla were enriched in one biome while others were enriched in another, indicating within-phylum differences in pathway content and habitat distributions. For instance, Fusobacteria pathways were predominantly host-enriched, but protein export, bacteria secretion system, and aminoacyl-tRNA biosynthesis were environment-enriched, indicating that these 3 pathways were more predominant in environment-enriched members of Fusobacteria. More generally, the two-component system, nitrogen metabolism, and galactose metabolism pathways were substantially more represented in the host-associated biome, while aminoacyl-tRNA biosynthesis was more predominant in the environmental biome (Figure 6B).

Biome-enrichment of CAZy families was restricted to fewer phyla than COG and KEGG groupings, likely reflecting a restriction of complex carbohydrate utilization to certain major clades (Figure 6C). For instance, most COG categories and KEGG pathways for Euryarchaeota and Synergistes were environment-enriched, but no CAZy families were enriched for either clade. Almost all of the highly abundant CAZy families belonging to Proteobacteria were environment-enriched, with the exception of GH04, GT01, GH43. Besides Proteobacteria, CAZy family enrichment in the environment was rather sparse, but some families were consistently enriched across many phyla, such as GT51, GT35, and GH13. Most CAZy families enriched in the host biome belonged to Firmicutes, Bacteroidetes, and Actinobacteria, with Firmicutes showing the highest diversity of enriched families. Notably, for most phyla, CAZy family enrichment was not exclusive to one biome, which likely reflects intra-phylum variation in CAZy family content for adaptation to each biome. For example, Firmicutes clusters annotated as GH57 and GH130 were environment-enriched, while all other Firmicutes CAZy families were host-enriched. In contrast, GT51 was predominantly environment-enriched, except for Firmicutes, indicating that CAZy enrichment can be contingent on the taxonomic and genomic context.

We performed the same method of mapping and enrichment testing for our multi-species metagenome data, with the goal of identifying functional groups enriched in Mammalia versus non-Mammalia. Compared to our host-environment analysis, we identified fewer significantly enriched features, which may be due to the smaller metagenome sample size or less pronounced partitioning of functional groups among biomes. For both Proteobacteria and Actinobacteria, most COG categories and KEGG pathways were enriched in non-mammals, with none enriched in Mammalia (Figure S15A & S15B). COG category enrichment in Mammalia was rather sparse, of which Spirochetes showed the highest number of categories enriched. KEGG pathway enrichment in Mammalia was much more robust, with many enriched pathways from the Spirochaetes, Bacteroidetes, Lentisphaerae, Fibrobacters, and Verrucomicrobia. Notably, many of the KEGG pathways differing between Mammalia and non-Mammilia were not the same as those differing between the host and environment biomes. For instance, bacterial secretion systems, streptomycin biosynthesis, ribosome, and protein export pathways significantly differed among the environment and host biomes, while quorum sensing, homologous recombination, and purine metabolism uniquely differentiated Mammalia versus non-Mammalia biomes (Figures 6 & S15).

*Functional metagenome profiling benefits from our gene catalogue*

We created a custom gene-level metagenome profiling database for the HUMAnN2 pipeline by merging our coding sequence catalogue with our previously constructed custom GTDBr89 database for HUMAnN2 (de la Cuesta-Zuluaga, Ley, and Youngblut 2019). We mapped our multi-species metagenomes to each database via the HUMAnN2 pipeline and compared the percent reads mapped. Due to the constraint of HUMAnN2 that all references must have a UniRef ID, we could only use 11.3% ($n$ = 722 795) of our gene clusters. Still, we found that including these clusters increased the mappability by 4 ± 5% s.d. (Figure S17). Mammalia species benefited the most, but at least one species from each class showed a mappability increase of >10% (Figure S17B).

### Supplemental Tables

**Table S1A.** Metadata for all samples in the multi-species metagenome dataset.

**Table S1B.** Summary of NCBI bioprojects used for the "multi-study" metagenome assemblies. "Number of samples used" indicates the number of metagenome samples used for metagenome assemblies. Datasets with >100 samples were randomly subsampled to 100. Metagenome assemblies were performed on a per-sample basis, with contig binning (generation of MAGs) performed on a per-study basis. Samples from BioProjects PRJNA316560 and PRJNA316570 were combined due to low numbers of samples and overlap in the animal host, methods, and study authors. There is no publication associated with BioProject PRJEB23642.

**Table S1C.** All samples obtained from the MGnify database in order to create the "host-environment" metagenome dataset.

**Table S1D.** Genome assembly accessions for all host species genomes used to filter out reads mapping to those genomes (the "vertebrata host genome" Kraken2 database shown in Figure 1).

**Table S1E.** Software parameters used.

**Table S2A.** Metadata on all non-redundant (de-replication at 99.9% ANI), quality (CheckM-estimated completeness ≥50% & contamination <5%) MAGs from all metagenome assemblies (those generated in this study and those published). Taxonomy and ANI values were derived from GTDB-Tk.

**Table S2B.** Metadata on all SGBs. Genome assembly metrics refer to the SGB reference genome.

**Table S2C.** Traitar-derived phenotype predictions for SGBs grouped by phylum and the biome for which they were enriched in (host, environment, or neither; see Figure 3C).

**Table S3A.** DESeq2 results for testing of host versus environment enrichment of SGB abundances (inferred via Krakenuniq). Only SGBs with a prevalence of >5% were included in the analysis (*n* = 967). Positive and negative $\log_2$ fold change values signify enrichment in host-associated and environment metagenomes, respectively. These data are shown in Figure 3.

**Table S3B.** DESeq2 results for testing of Mammalia versus non-Mammalia enrichment of SGB abundances (inferred via Krakenuniq). Only SGBs with a prevalence of >5% were included in the analysis (*n* = 663). Positive and negative $\log_2$ fold change values signify enrichment in Mammalia and non-Mammalia metagenomes, respectively. These data are shown in Figure S8.

**Table S4A.** All SGB BGCs identified by AntiSMASH and clustered by BiGSCAPE into clans and families. These data are shown in Figure S10.

**Table S4B.** All AMR markers identified by Abricate, summed by species. These data are shown in Figure S9.

**Table S5A.** DESeq2 results for testing of host versus environment enrichment of gene cluster abundances (50% sequence identity clustering) summed by COG functional category and phylum. Positive and negative $\log_2$ fold change values signify enrichment in host-associated and environment metagenomes, respectively. These data are shown in Figure 6.

**Table S5B.** DESeq2 results for testing of host versus environment enrichment of gene cluster abundances (50% sequence identity clustering) summed by KEGG pathway and phylum. Positive and negative $\log_2$ fold change values signify enrichment in host-associated and environment metagenomes, respectively. These data are shown in Figure 6.

**Table S5C.** DESeq2 results for testing of host versus environment enrichment of gene cluster abundances (50% sequence identity clustering) summed by CAZy family and phylum. Positive and negative $\log_2$ fold change values signify enrichment in host-associated and environment metagenomes, respectively. These data are shown in Figure 6.

**Table S5D.** DESeq2 results for testing of Mammalia versus non-Mammalia enrichment of gene cluster abundances (50% sequence identity clustering) summed by COG functional category and phylum. Positive and negative $\log_2$ fold change values signify enrichment in Mammalia and non-Mammalia metagenomes, respectively. These data are shown in Figure S15.

**Table S5E.** DESeq2 results for testing of Mammalia versus non-Mammalia enrichment of gene cluster abundances (50% sequence identity clustering) summed by KEGG pathway and phylum. Positive and negative $\log_2$ fold change values signify enrichment in Mammalia and non-Mammalia metagenomes, respectively. These data are shown in Figure S15.

**Table S5F.** DESeq2 results for testing of Mammalia versus non-Mammalia enrichment of gene cluster abundances (50% sequence identity clustering) summed by CAZy family and phylum. Positive and negative $\log_2$ fold change values signify enrichment in Mammalia and non-Mammalia metagenomes, respectively. These data are shown in Figure S15.

### Supplemental Figures

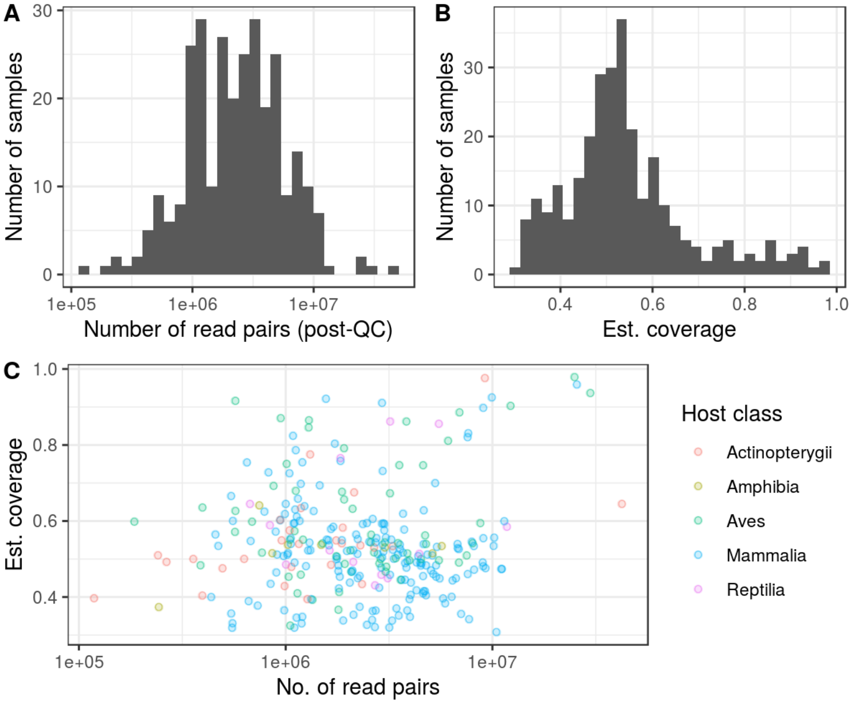


**Figure S1.** Read quality control (QC) of the “multi-species” animal gut metagenome dataset generated in this study. A) The number of post-QC Illumina HiSeq read pairs per sample. B) The per-sample nonpareil-estimated coverage of diversity (a cutoff of >0.3 used to filter samples). C) No strong associations between the number of paired-end reads, estimated coverage, and host taxonomic class.

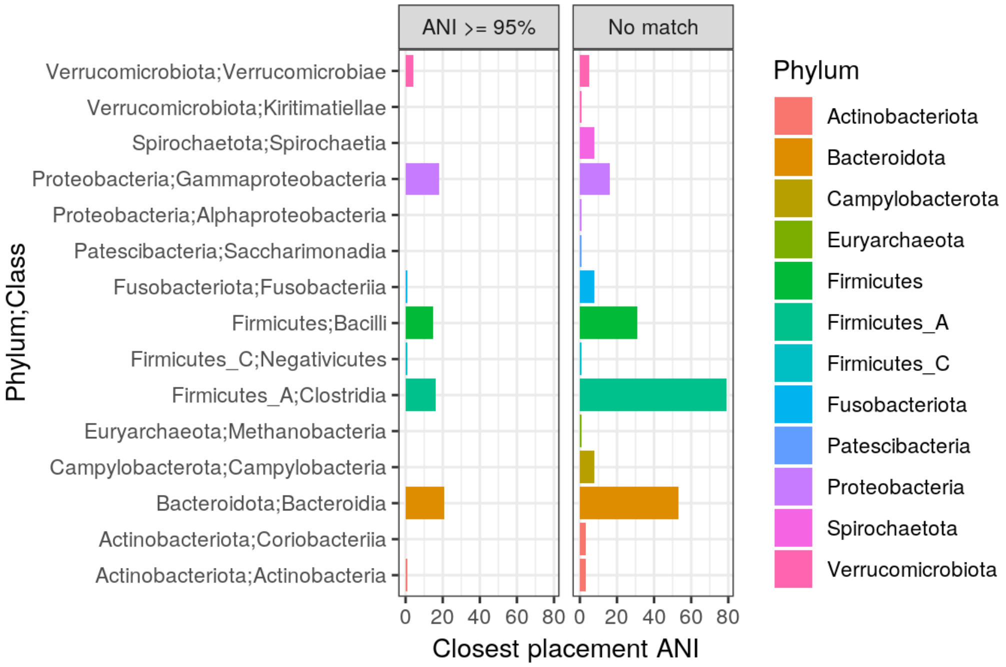


**Figure S2.** The number of quality MAGs (99.9% ANI de-replicated) per phylum and class, which were generated from the multi-species dataset metagenome assemblies. The plot is faceted by MAGs with a species-level hit to the GTDB (“ANI >= 95%”) or not (“No match”).

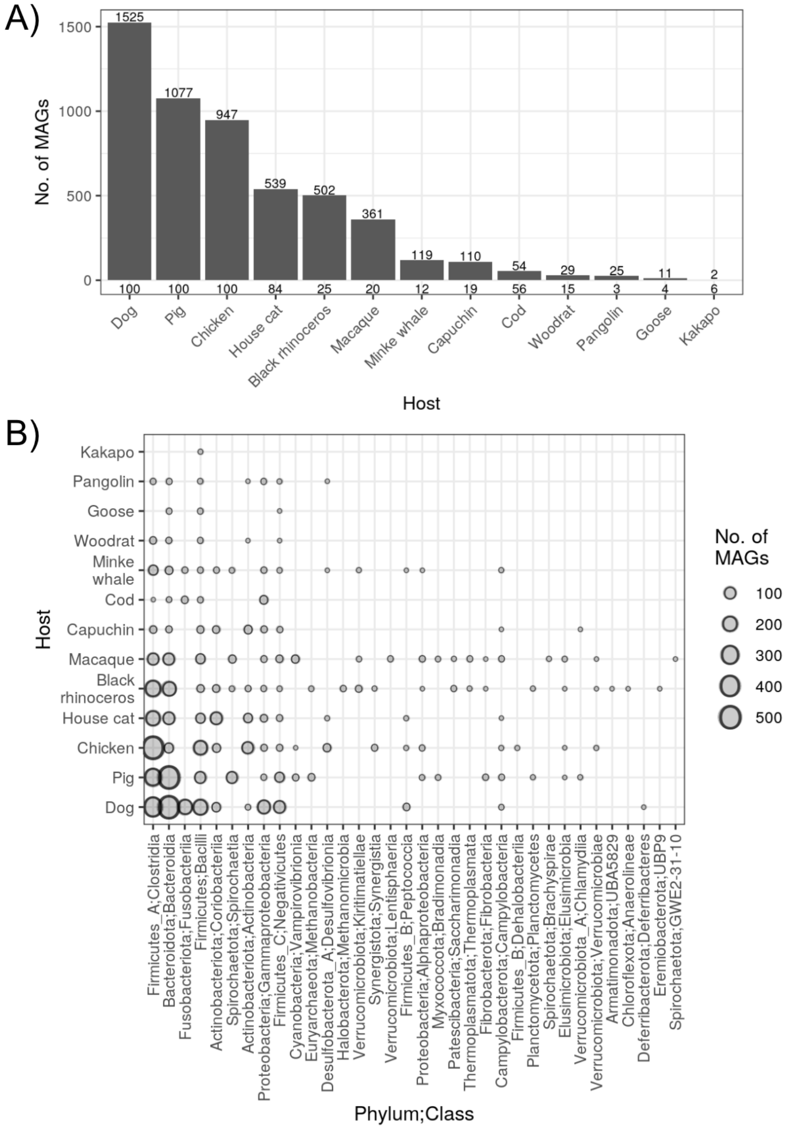


**Figure S3.** A) The number of quality MAGs per metagenome dataset. The value above each bar is the number of MAGs, and the value below each bar is the number of samples used for each metagenome assembly pipeline run. Note that the “Goose” gut metagenomes comprise 2 studies (see Table S1B). B) The number of quality MAGs per dataset, grouped by phylum and class.

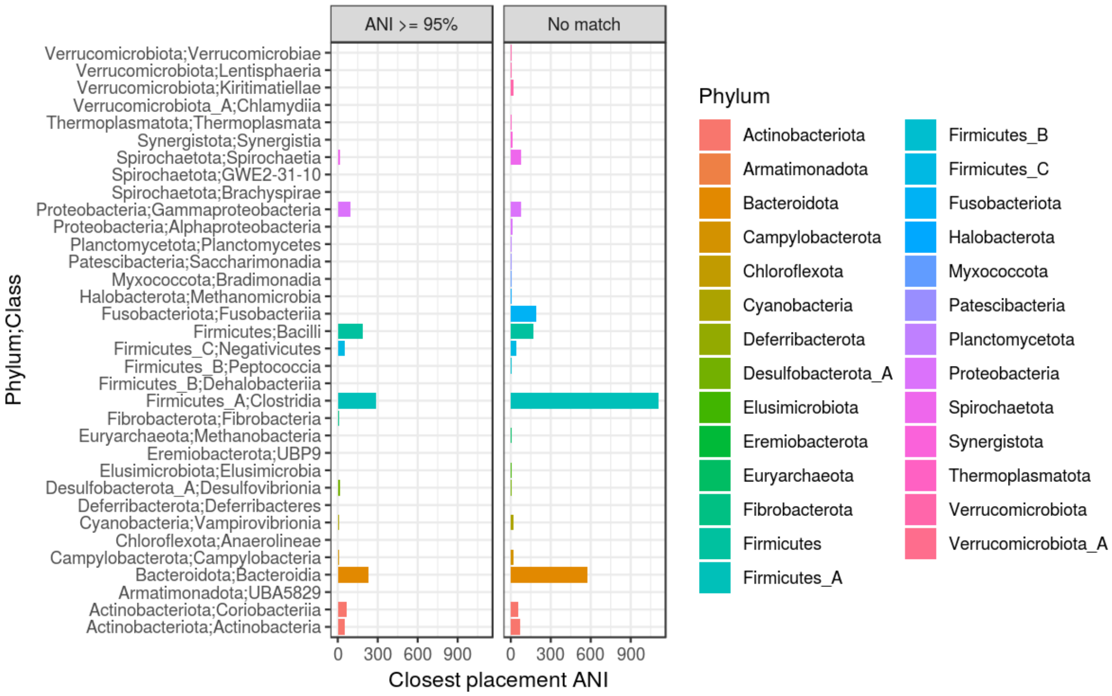


**Figure S4.** The number of quality MAGs (99.9% ANI de-replicated) per phylum and class. The MAGs are all those generated from the multi-species and published dataset assemblies. The plot is faceted by MAGs with a species-level hit to the GTDB (“ANI >= 95%”) or not (“No match”).

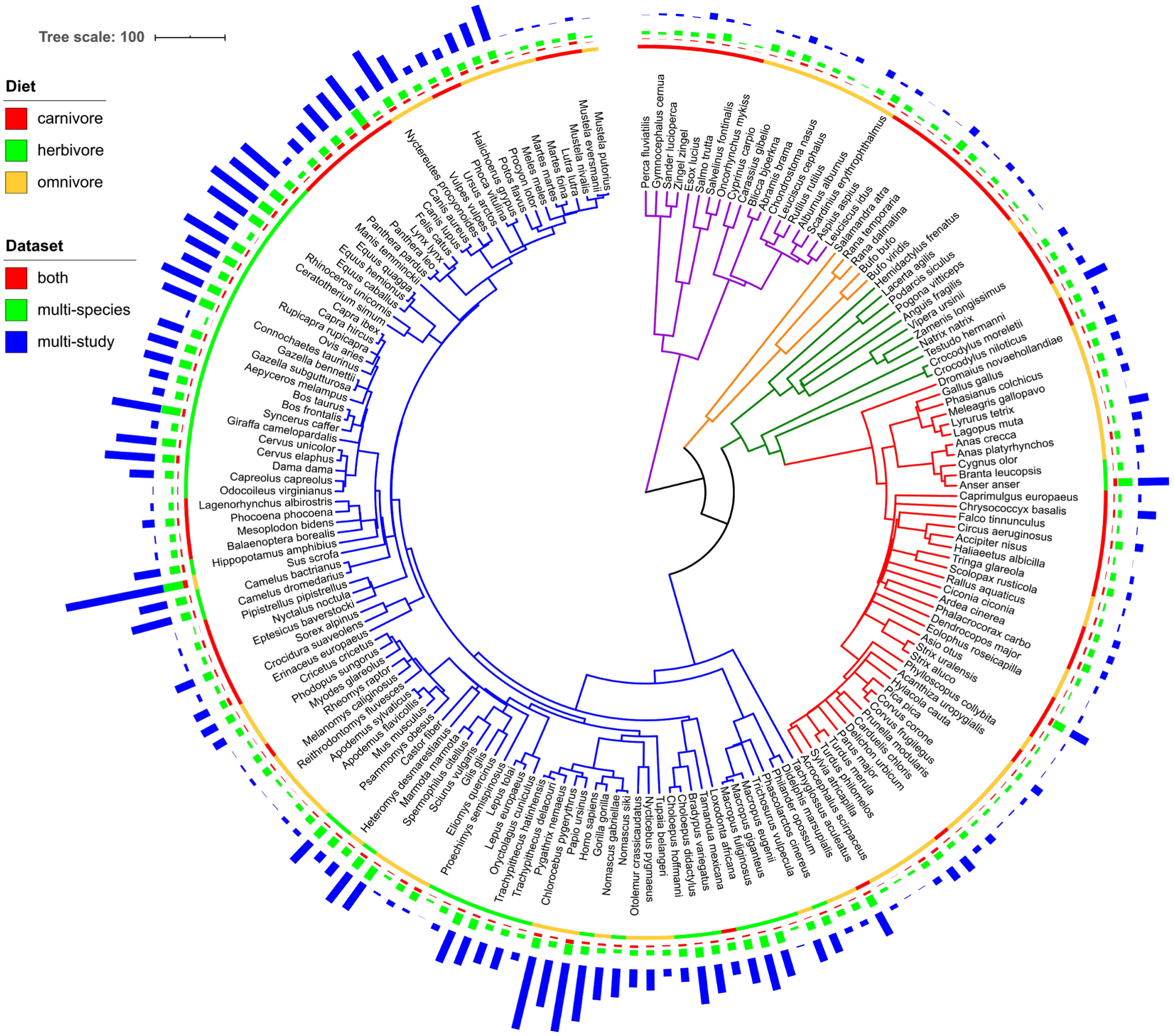


**Figure S5.** *The SGB collection is representative of taxa found across vertebrates.* The colored bar charts show the number of SGBs present in each sample as determined by mapping metagenome reads to each SGB representative genome via Krakenuniq. The bars are colored by the dataset of origin ("multi-species" or "multi-study", see Methods) for each SGB, with SGBs labeled as "both" if the SGB encompassed MAGs assembled from both the multi-species and multi-study datasets. The minimum and maximum bar sizes represent 1 and 532 SGBs, respectively. The tree is the same as shown in Figure 1.

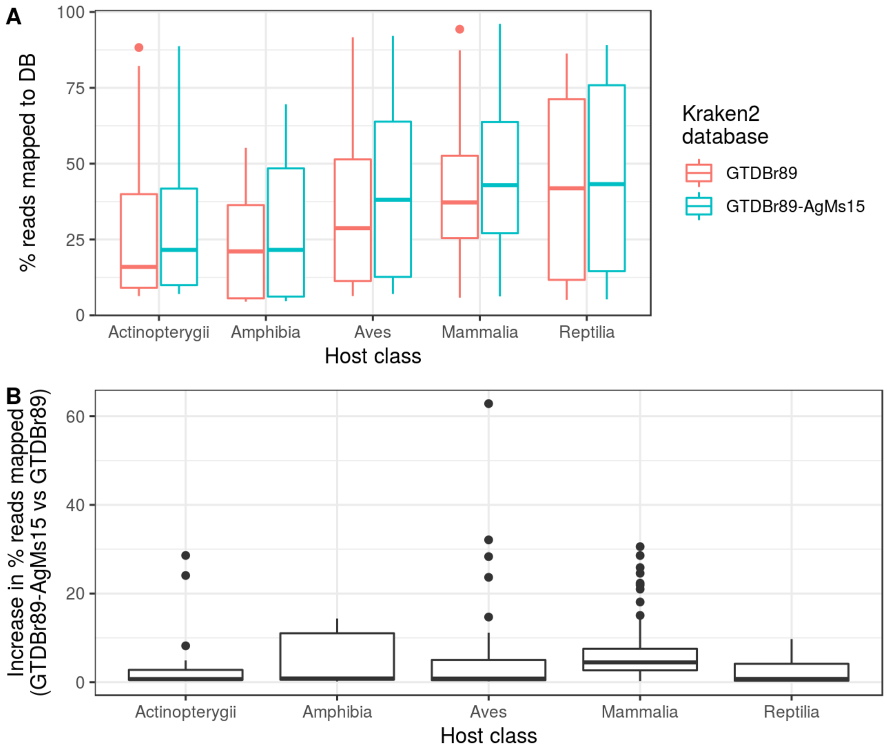


**Figure S6.** A) The percent of reads from our multi-species dataset mapped via Kraken2 to either the custom GTDB Release-89 database ("GTDBr89") or the same database with all SGBs from this study included ("GTDBr89-AgMs15"). The increase in percent reads mapped was significant based on a one-sided paired t-test comparing all GTDBr89 versus GTDBr89-AgMs15 samples ($P < 0.005$). B) Percent increase of reads mapped when using the GTDBr89-AgMs15 database versus GTDBr89. Boxplot centerlines, edges, whiskers, and points signify the median, interquartile range (IQR), 1.5× IQR, and >1.5× IQR, respectively.

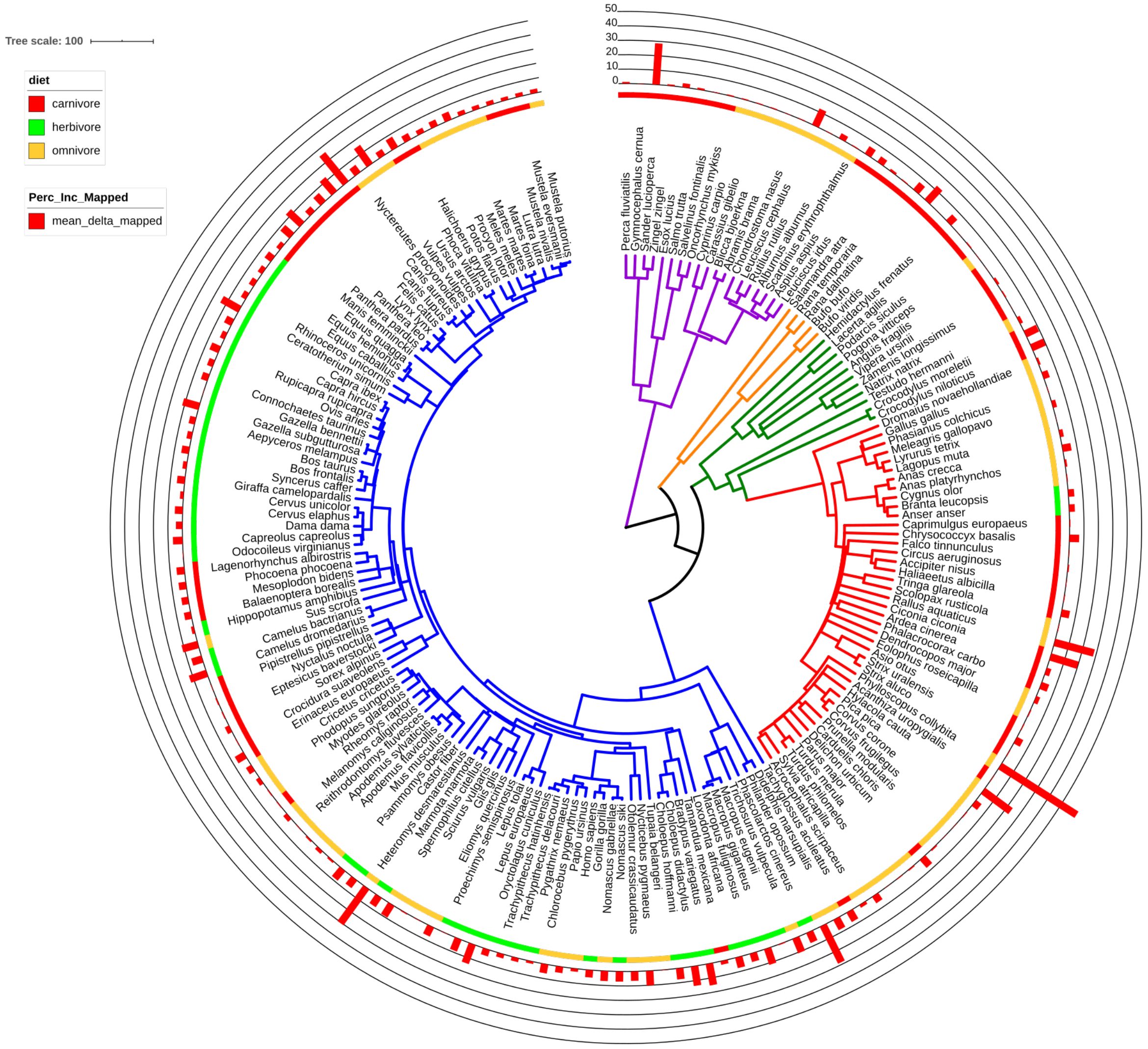


**Figure S7.** Mean percent increase ("mean_delta_mapped") in mapped reads for each metagenome sample when using the GTDBr89-AgMs15 reference database versus GTDB-r89 (see Figure S6). The tree is the same as in Figure 1.

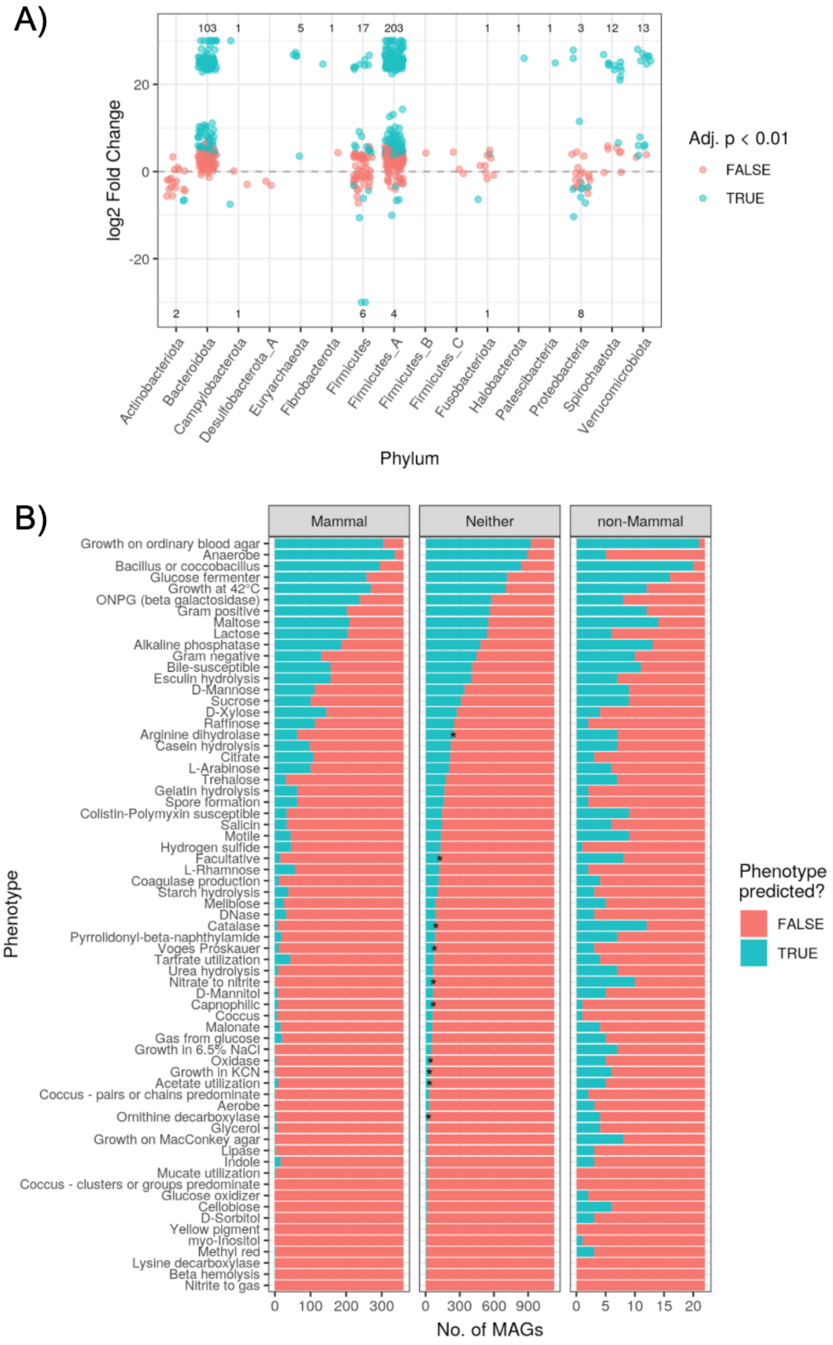


**Figure S8.** A) The number of SGBs significantly enriched in Mammalia (positive log2 fold change; “l2fc”) or non-Mammalia gut metagenomes (negative l2fc). Values shown are the number of MAGs significantly enriched (blue) in either biome or not found to be significant (red). B) No signal of distinct phenotypes among SGBs enriched in Mammal or non-Mammal gut metagenomes. Predicted phenotypes are summarized for the SGBs significantly enriched (DESeq2, *adj. P* < 0.01) in Mammal or non-Mammal metagenomes (x-axis facet). Note the difference in x-axis scale. Asterisks denote phenotypes significantly more prevalent in SGBs of the particular biome versus a null model of 1000 permutations in which biome labels were shuffled among SGBs. See Table S3B for all DESeq2 results.

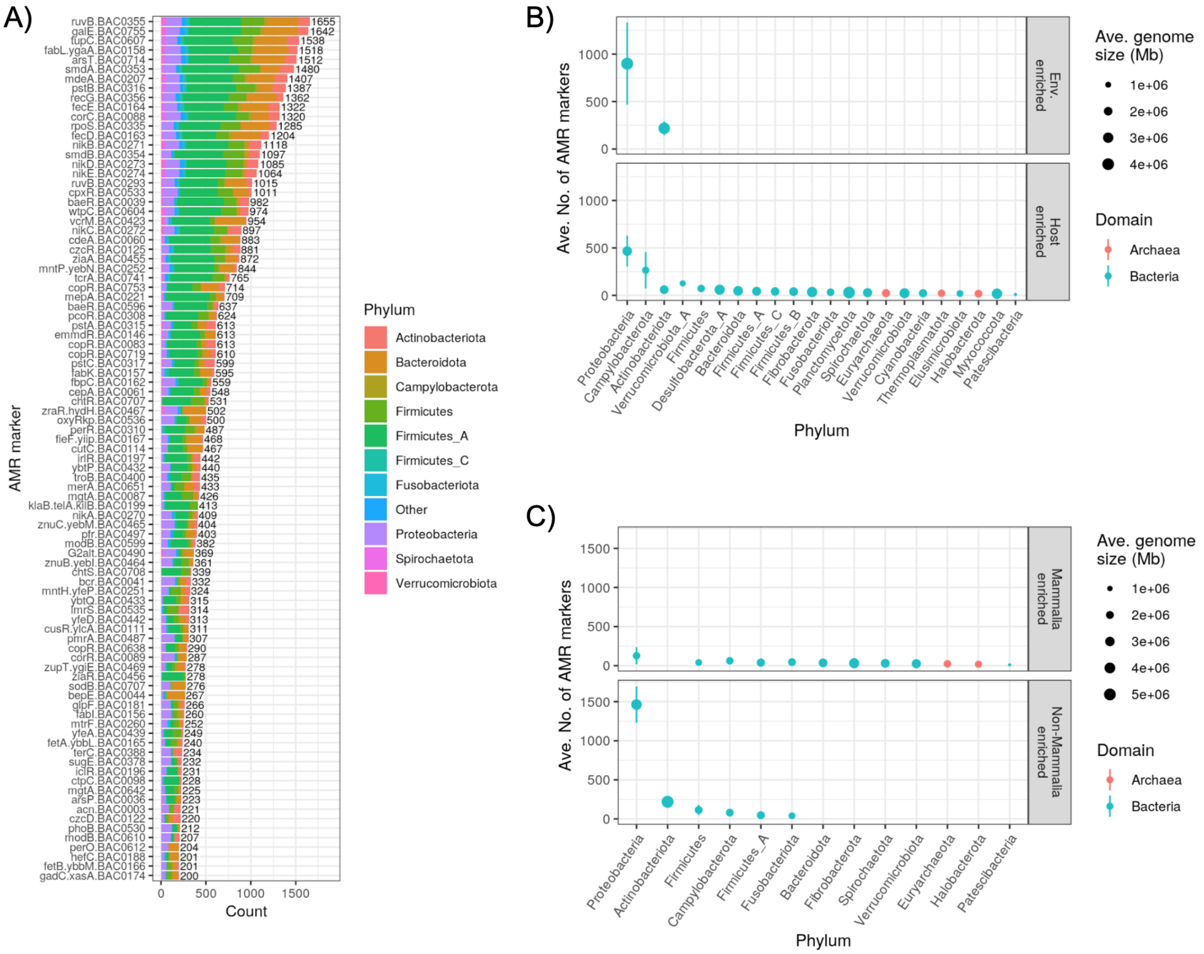


**Figure S9.** A) The number of genes per AMR marker and phylum. For clarity, only markers with ≥200 total genes are shown, and all phyla with <400 AMR genes are grouped as "Other". The average number of antimicrobial resistance (AMR) genes per genome in SGBs enriched in B) host or environmental metagenomes C) Mammalia versus non-Mammalia gut metagenomes. Points represent mean among all SGBs in each phylum, and line ranges denote the standard error of the mean.

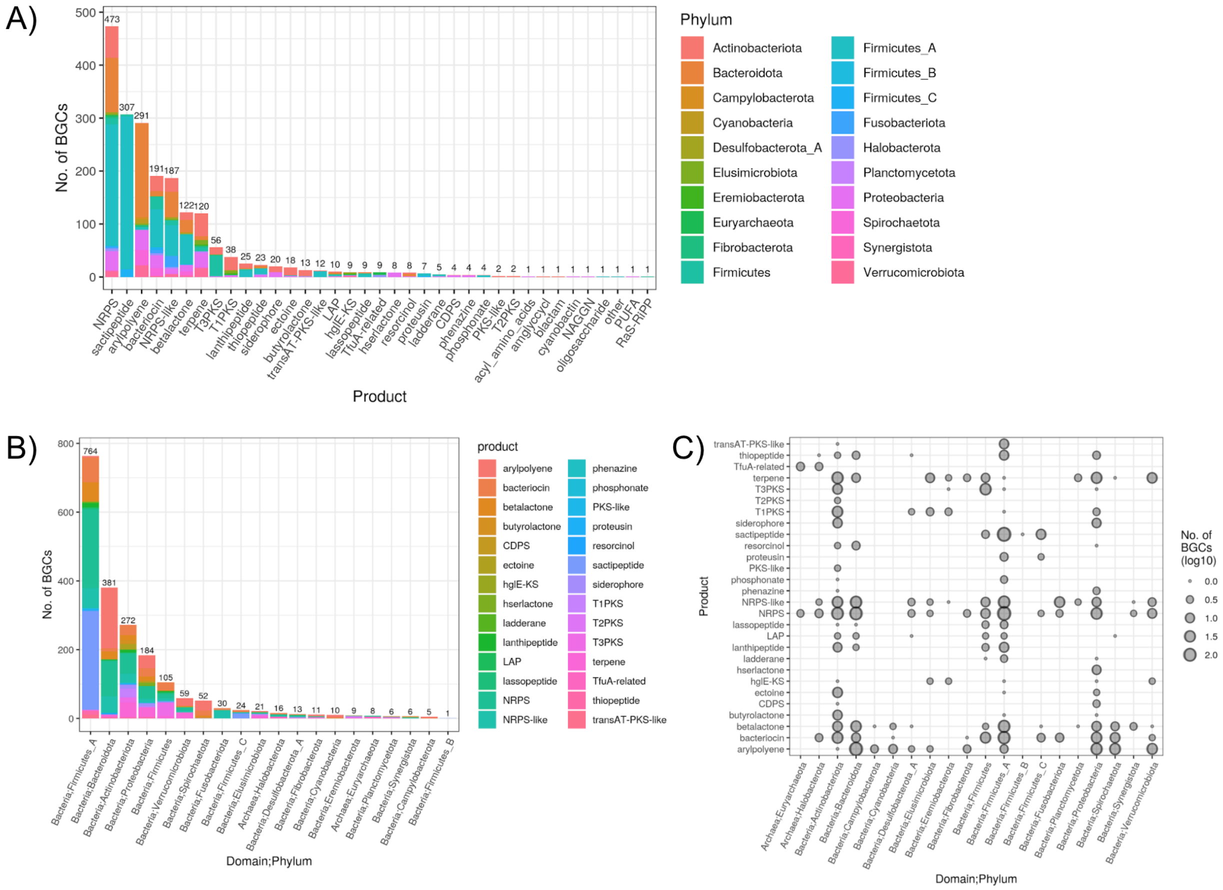


**Figure S10.** The number of SGB BGCs identified by AntiSMASH grouped by A) BGC product (x-axis) and Phylum (color) or B) *vice versa*. The plot in C) is an alternative view of A) and B). See Table S4A for a list of all BGCs.

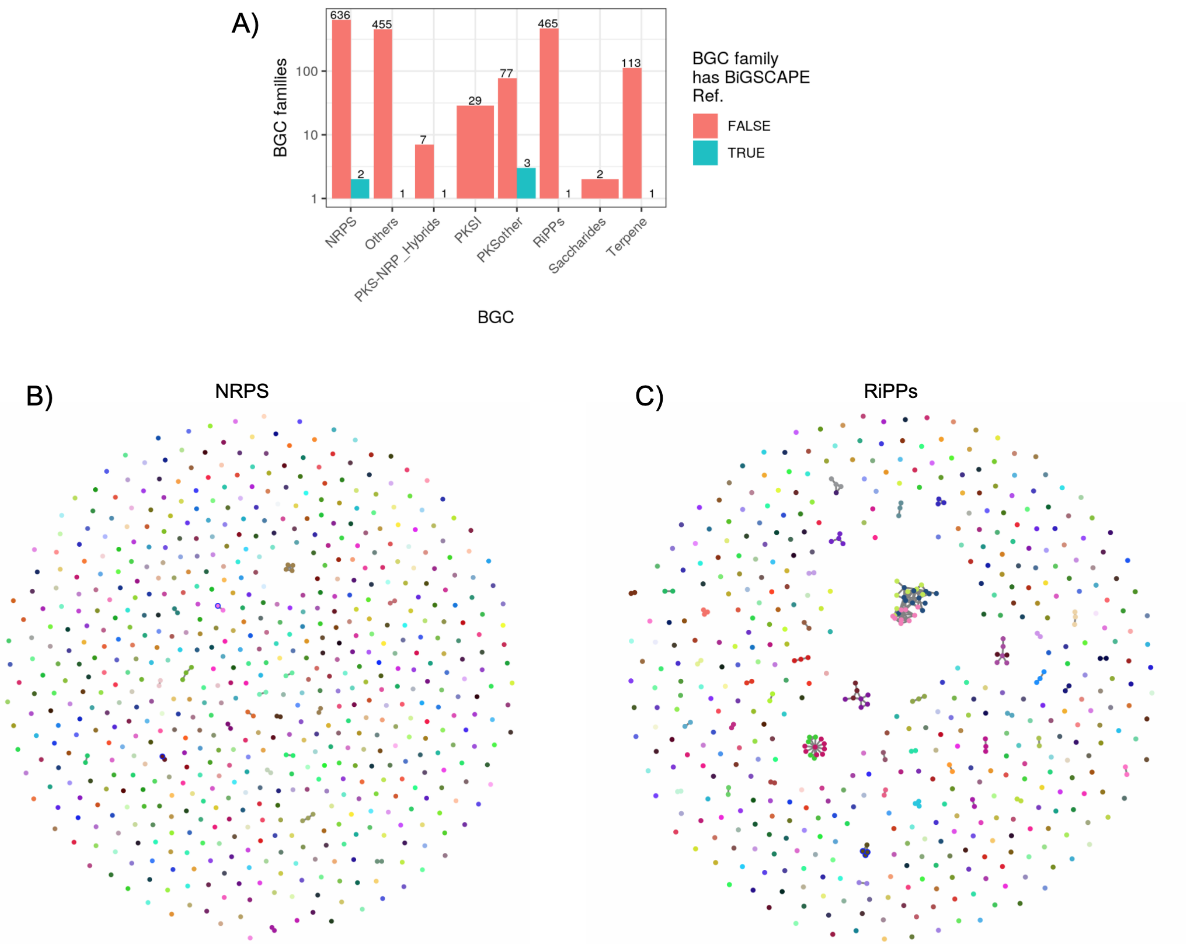


**Figure S11.** A) The number of BGC families from all SGB BGCs which have or do not include a BiGSCAPE reference, grouped by BiGSCAPE class. The BGC families are plotted by BGC product type, with “Others” representing all other product types (see Figure S10 for a full list of products). The networks in B) and C) show BGC similarity of the two largest BGC classes: NRPS and RiPPS. The nodes represent individual BGCs, and edges connect BGCs in the same family. See Table S4B for a list of all AMR markers.

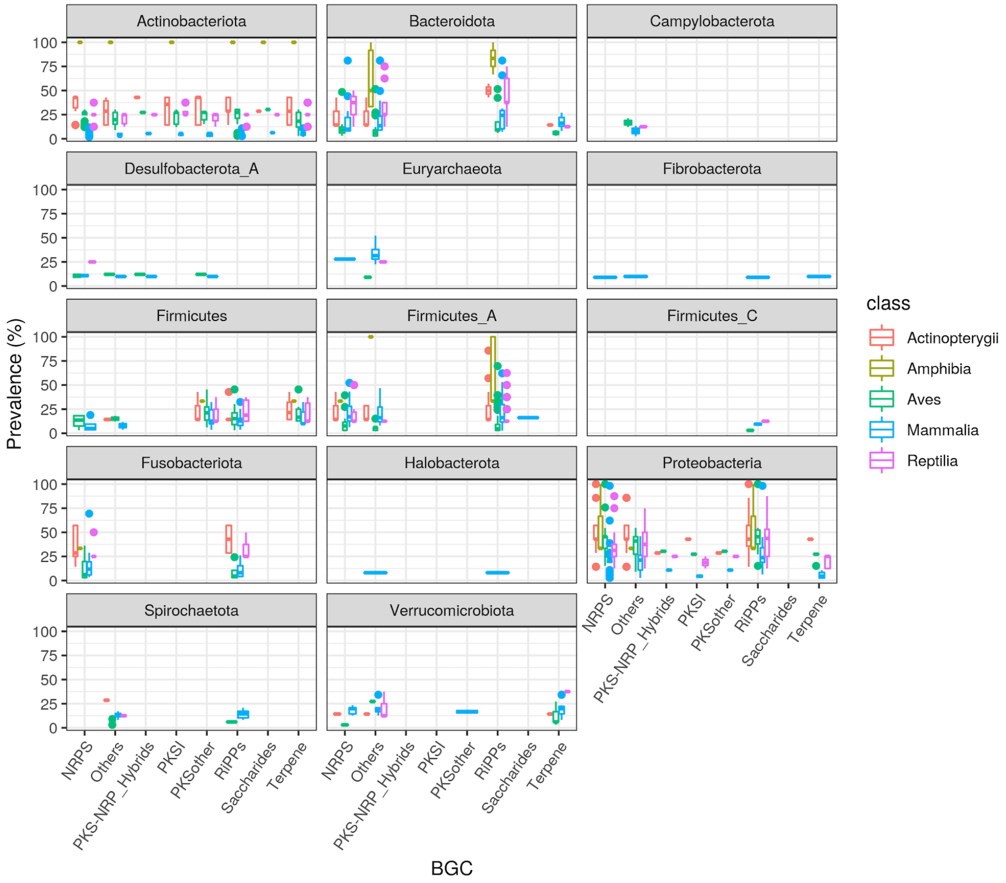


**Figure S12.** The prevalence (*i.e.,* the percent of samples where observed) of all SGB BGCs identified by AntiSMASH across all metagenomes in our multi-species dataset. Boxplot centerlines, edges, whiskers, and points signify the median, interquartile range (IQR), 1.5× IQR, and >1.5× IQR, respectively.

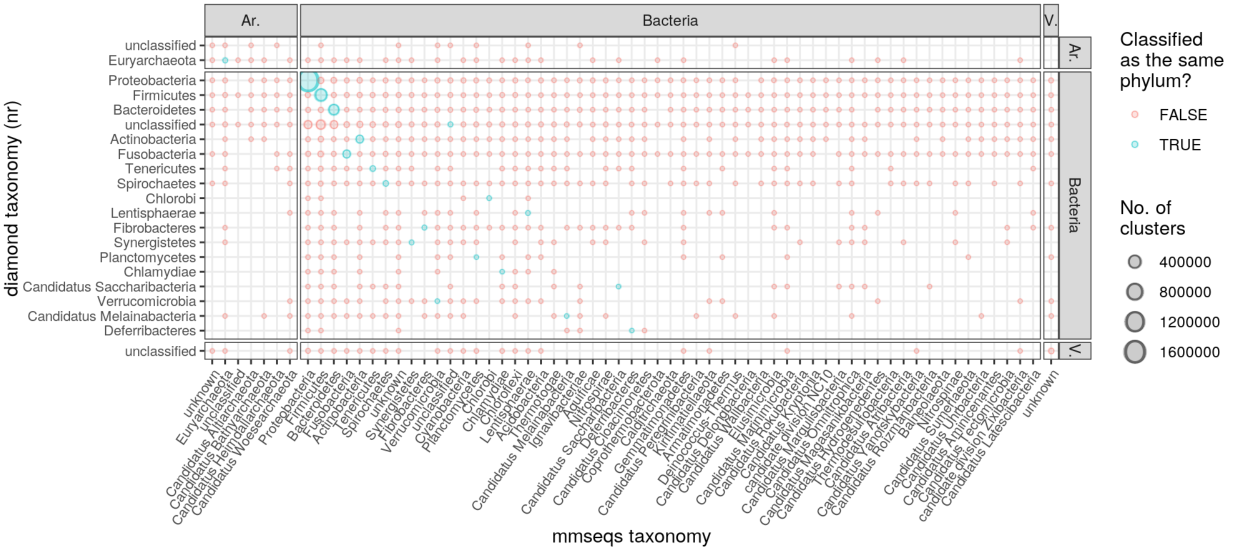


**Figure S13**. A comparison of mmseqs versus DIAMOND (NCBI nr database) taxonomic classification of archaeal ("Ar."), bacterial ("Bacteria"), and viral ("V.") clusters. The size of the points represents the number of gene clusters with the particular phylum-level taxonomic classification. For clarity, only phyla with ≥100 clusters (50% sequence identity clustering) are shown. The total number of archaeal, bacterial, and viral clusters classified by mmseqs2 and DIAMOND were 3,079,918 and 3,146,914, respectively.

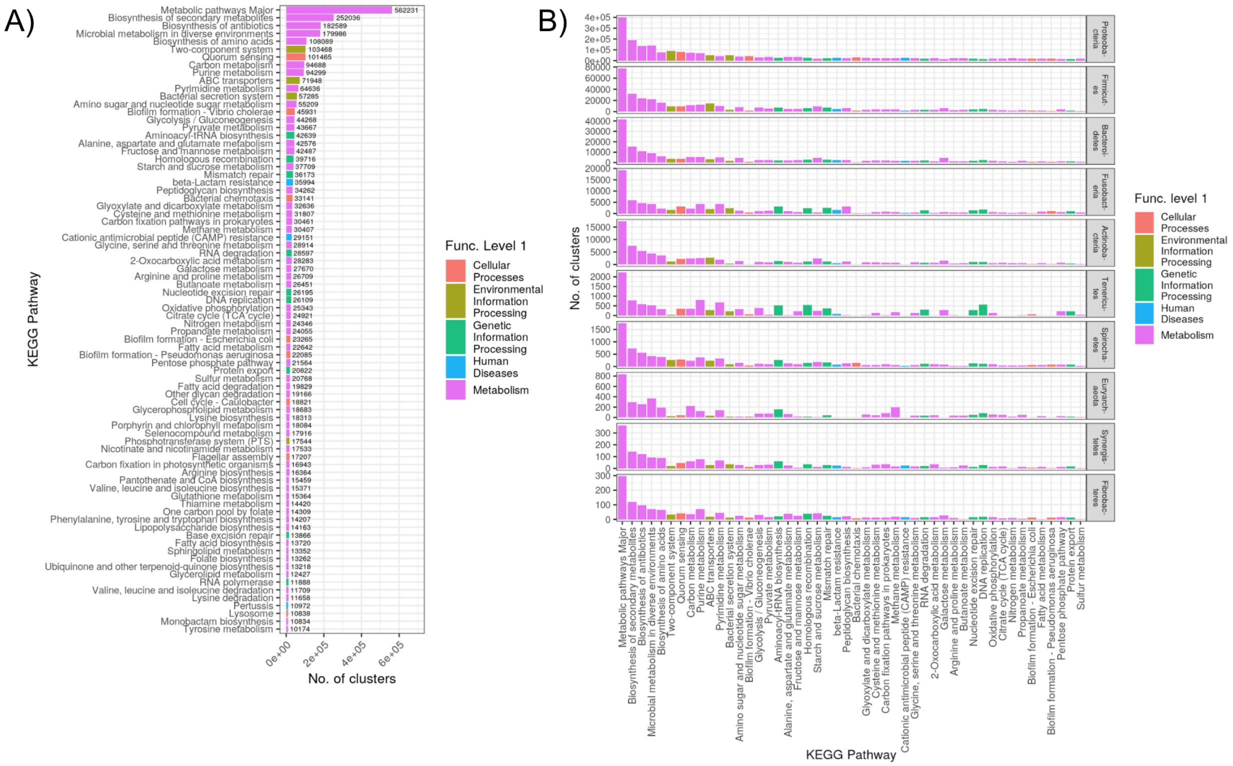


**Figure S14**. A) The number of clusters associated with each KEGG pathway, colored by KEGG functional level 1. For clarity, only pathways with ≥10,000 clusters are shown. B) The number of clusters associated with each KEGG pathway, broken down by phylum and color by KEGG functional level 1. Note the different y-axis scales. For clarity, only phyla with ≥5,000 clusters and only KEGG pathways with ≥10,000 clusters are shown.

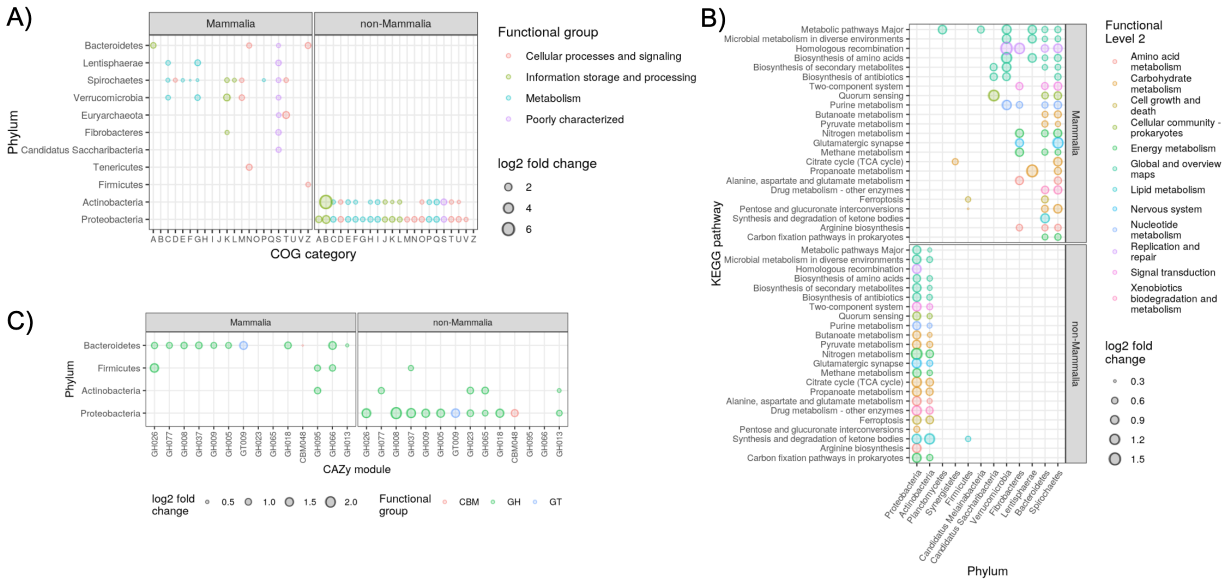


**Figure S15**. Enrichment of gene clusters grouped by phylum and A) COG category B) KEGG pathway or C) CAZy family. Only grouping significantly enriched in abundance (DESeq2, *adj. P* < 1e-5) in either biome are shown. Only gene clusters observed in at least 25% of the metagenomes were included. For clarity, only KEGG pathways enriched in >7 phyla are shown, and only CAZy families enriched in >1 phylum are shown. Note that the axes are flipped in B) relative to A) and C). See Tables S5D, S5E, and S5F for all DESeq2 results.

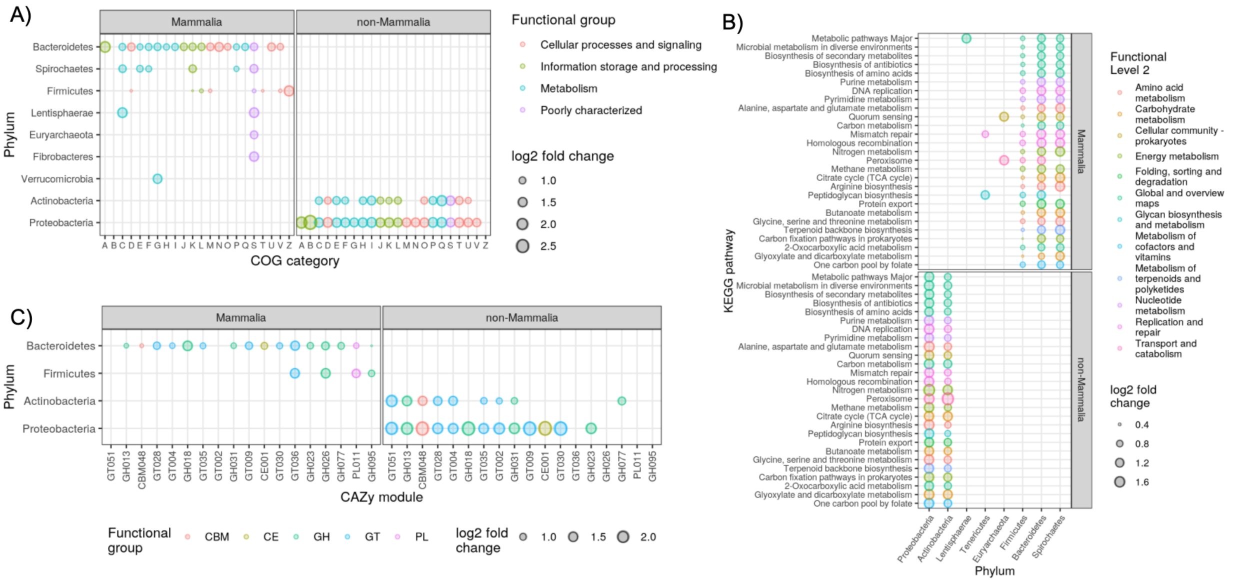


**Figure S16.** The same as Figure S15, but more strict DIAMOND search parameters, with a sequence identity cutoff of 50% instead of no such cutoff. The stricter parameters substantially reduced the number of gene clusters observed in any metagenome (48% fewer clusters), but the results are qualitatively similar to Figure S15.

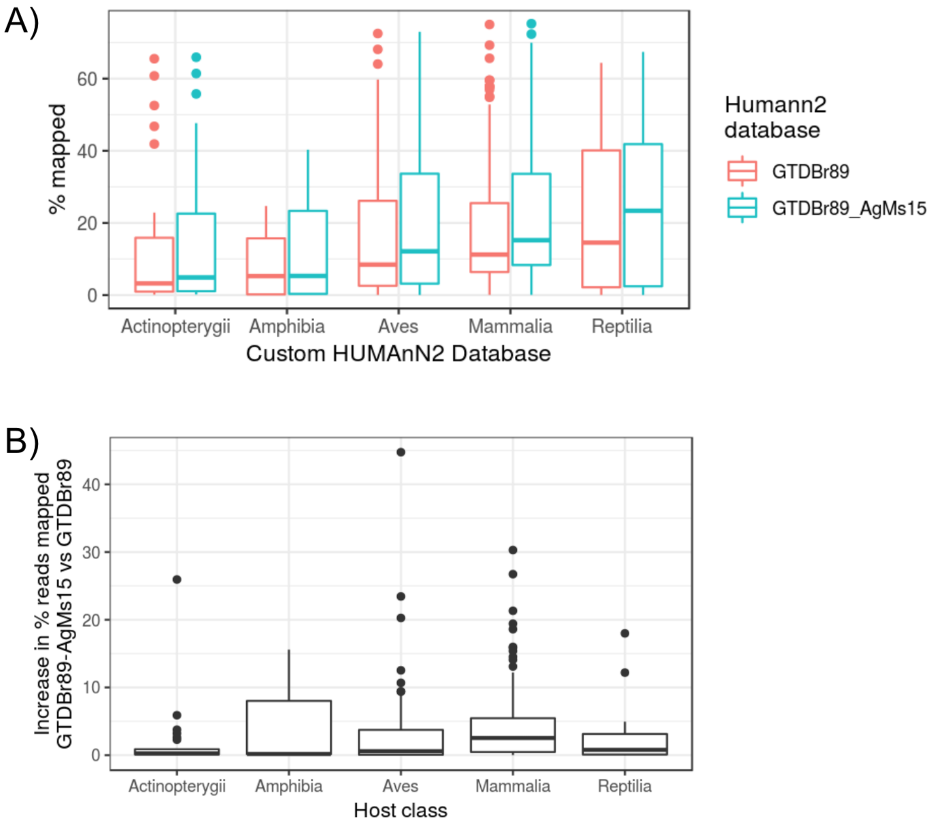


**Figure S17.** A) The percent of reads from our multi-species metagenome dataset mapped via the HUMAnN2 pipeline to either the custom GTDB Release-89 database ("GTDBr89") or the same database with all 50% sequence identity gene clusters generated from our gene-based metagenome assemblies ("GTDBr89-AgMs15"). B) The percent increase of reads mapped per metagenome sample (grouped by class) when using the GTDBr89-AgMs15 database versus GTDBr89. Boxplot centerlines, edges, whiskers, and points signify the median, interquartile range (IQR), 1.5× IQR, and >1.5× IQR, respectively.

Jiang, Hongshan, Rong Lei, Shou-Wei Ding, and Shuifang Zhu. 2014. “Skewer: A Fast and Accurate Adapter Trimmer for next-Generation Sequencing Paired-End Reads.” *BMC Bioinformatics* 15 (June): 182.

Karasov, Talia L., Juliana Almario, Claudia Friedemann, Wei Ding, Michael Giolai, Darren Heavens, Sonja Kersten, et al. 2018. “Arabidopsis Thaliana and Pseudomonas Pathogens Exhibit Stable Associations over Evolutionary Timescales.” *Cell Host & Microbe* 24 (1): 168–79.e4.

Kautsar, Satria A., Kai Blin, Simon Shaw, Jorge C. Navarro-Muñoz, Barbara R. Terlouw, Justin J. J. van der Hooft, Jeffrey A. van Santen, et al. 2020. “MIBiG 2.0: A Repository for Biosynthetic Gene Clusters of Known Function.” *Nucleic Acids Research* 48 (D1): D454–58.

Köster, Johannes, and Sven Rahmann. 2012. “Snakemake--a Scalable Bioinformatics Workflow Engine.” *Bioinformatics* 28 (19): 2520–22.

Langmead, Ben, and Steven L. Salzberg. 2012. “Fast Gapped-Read Alignment with Bowtie 2.” *Nature Methods* 9 (4): 357–59.

Love, Michael I., Wolfgang Huber, and Simon Anders. 2014. “Moderated Estimation of Fold Change and Dispersion for RNA-Seq Data with DESeq2.” *Genome Biology* 15 (12): 550.

Lu, Jennifer, Florian P. Breitwieser, Peter Thielen, and Steven L. Salzberg. 2017. “Bracken: Estimating Species Abundance in Metagenomics Data.” *PeerJ Computer Science* 3 (January): e104.

Meyer, Fernando, Peter Hofmann, Peter Belmann, Ruben Garrido-Oter, Adrian Fritz, Alexander Sczyrba, and Alice C. McHardy. 2018. “AMBER: Assessment of Metagenome BinnERs.” *GigaScience* 7 (6). https://doi.org/10.1093/gigascience/giy069.

Mitchell, Alex L., Alexandre Almeida, Martin Beracochea, Miguel Boland, Josephine Burgin, Guy Cochrane, Michael R. Crusoe, et al. 2020. “MGnify: The Microbiome Analysis Resource in 2020.” *Nucleic Acids Research* 48 (D1): D570–78.

Navarro-Muñoz, Jorge C., Nelly Selem-Mojica, Michael W. Mullowney, Satria A. Kautsar, James H. Tryon, Elizabeth I. Parkinson, Emmanuel L. C. De Los Santos, et al. 2020. “A Computational Framework to Explore Large-Scale Biosynthetic Diversity.” *Nature Chemical Biology* 16 (1): 60–68.

Nurk, Sergey, Dmitry Meleshko, Anton Korobeynikov, and Pavel A. Pevzner. 2017. “metaSPAdes: A New Versatile Metagenomic Assembler.” *Genome Research* 27 (5): 824–34.

Olm, Matthew R., Alexander Crits-Christoph, Spencer Diamond, Adi Lavy, Paula B. Matheus Carnevali, and Jillian F. Banfield. 2020. “Consistent Metagenome-Derived Metrics Verify and Delineate Bacterial Species Boundaries.” *mSystems* 5 (1). https://doi.org/10.1128/mSystems.00731-19.

Parks, Donovan H., Maria Chuvochina, David W. Waite, Christian Rinke, Adam Skarshewski, Pierre-Alain Chaumeil, and Philip Hugenholtz. 2018. “A Standardized Bacterial Taxonomy Based on Genome Phylogeny Substantially Revises the Tree of Life.” *Nature Biotechnology* 36 (10): 996–1004.

Parks, Donovan H., Michael Imelfort, Connor T. Skennerton, Philip Hugenholtz, and Gene W. Tyson. 2015. “CheckM: Assessing the Quality of Microbial Genomes Recovered from Isolates, Single Cells, and Metagenomes.” *Genome Research* 25 (7): 1043–55.

Pierce, N. Tessa, Luiz Irber, Taylor Reiter, Phillip Brooks, and C. Titus Brown. 2019. “Large-Scale Sequence Comparisons with Sourmash.” *F1000Research* 8 (July): 1006.

R Core Team. 2020. *R: A Language and Environment for Statistical Computing*. Vienna, Austria: R Foundation for Statistical Computing.

Rodriguez-R, Luis M., Santosh Gunturu, James M. Tiedje, James R. Cole, and Konstantinos T. Konstantinidis. 2018. “Nonpareil 3: Fast Estimation of Metagenomic Coverage and Sequence Diversity.” *mSystems* 3 (3). https://doi.org/10.1128/mSystems.00039-18.

Segata, Nicola, Daniela Börnigen, Xochitl C. Morgan, and Curtis Huttenhower. 2013. “PhyloPhlAn Is a New Method for Improved Phylogenetic and Taxonomic Placement of Microbes.” *Nature Communications* 4: 2304.

Sieber, Christian M. K., Alexander J. Probst, Allison Sharrar, Brian C. Thomas, Matthias Hess, Susannah G. Tringe, and Jillian F. Banfield. 2018. “Recovery of Genomes from Metagenomes via a Dereplication, Aggregation and Scoring Strategy.” *Nature Microbiology* 3 (7): 836–43.

Steinegger, Martin, Milot Mirdita, and Johannes Söding. 2019. “Protein-Level Assembly Increases Protein Sequence Recovery from Metagenomic Samples Manyfold.” *Nature Methods* 16 (7): 603–6.

Steinegger, Martin, and Johannes Söding. 2018. “Clustering Huge Protein Sequence Sets in Linear Time.” *Nature Communications* 9 (1): 2542.

Vollmers, John, Sandra Wiegand, and Anne-Kristin Kaster. 2017. “Comparing and Evaluating Metagenome Assembly Tools from a Microbiologist’s Perspective - Not Only Size Matters!” *PloS One* 12 (1): e0169662.

Walker, Bruce J., Thomas Abeel, Terrance Shea, Margaret Priest, Amr Abouelliel, Sharadha Sakthikumar, Christina A. Cuomo, et al. 2014. “Pilon: An Integrated Tool for Comprehensive Microbial Variant Detection and Genome Assembly Improvement.” *PloS One* 9 (11): e112963.

Wickham, Hadley. 2009. *ggplot2: Elegant Graphics for Data Analysis*. 1st ed. 2009. Corr. 3rd printing 2010 edition. New York: Springer. 2016. “Tidyr: Easily Tidy Data with `spread()` and `gather()` Functions.” https://CRAN.R-project.org/package=tidyr.

Wickham, Hadley, and Jennifer Bryan. 2019. “Readxl: Read Excel Files.” https://CRAN.R-project.org/package=readxl.

Wickham, Hadley, and Romain Francois. 2016. “Dplyr: A Grammar of Data Manipulation.” https://CRAN.R-project.org/package=dplyr.

Youngblut, Nicholas D., Georg H. Reischer, William Walters, Nathalie Schuster, Chris Walzer, Gabrielle Stalder, Ruth E. Ley, and Andreas H. Farnleitner. 2019. “Host Diet and Evolutionary History Explain Different Aspects of Gut Microbiome Diversity among Vertebrate Clades.” *Nature Communications* 10 (1): 2200.
